# Algorithmic dissection of optic flow memory in larval zebrafish

**DOI:** 10.1101/2025.04.15.648832

**Authors:** Ryosuke Tanaka, Ruben Portugues

## Abstract

The visual stabilization behavior in the larval zebrafish reflects the history of optic flow experienced in the recent past. This integrative process has gained traction in recent years as a simplified, tractable model of working memory and decision-making. Yet, its algorithmic bases are poorly understood. In the present study, we first demonstrate that fish integrate externally generated optic flow over time, disregarding self-generated optic flow. This observation suggests that the fish integrates optic flow to better estimate the state of the environment, rather than to keep track of their position. Second, through reverse correlation and delay-based paradigms, we reveal multiple timescales involved in the optic flow integration. With the help of quantitative modeling, we show that the fish becomes more forgetful about the past optic flow in a more dynamic visual environment. Next, with whole-brain, light-sheet calcium imaging, we find that optic flow selective neurons throughout the brain exhibit neural signatures of motor efference copies, mirroring the behavioral findings. Lastly, with two photon calcium imaging, we show that inferior olive neurons integrate forward and backward flow separately, giving clues to how the multiple timescales of the optic flow integration is implemented. Overall, the results here refine our algorithmic and functional understanding of the history dependence of the visual stabilization behaviors in the larval zebrafish, paving the way for deciphering its circuit implementations.

## Introduction

The behaviors of animals reflect not only the incoming sensory inputs but also the history of sensory events. How a brain, which is composed of neurons operating at the timescale of milliseconds, can retain information over a longer, seconds timescale is not trivial, and has attracted researchers’ attention [1–3]. A relatively simple example of such history-dependent behaviors is the optomotor response in the larval zebrafish – one of the smallest vertebrate model organisms in the modern neuroscience. When presented with patterns of wide-field visual motion indicative of observer movements (i.e. optic flow [4]), various animal species including the larval zebrafish exhibit body movements to stabilize their heading and position, a behavior called optomotor response (OMR) [5–8]. Recently, it has been shown that the OMR of the larval zebrafish can be affected by optic flow that happened many seconds ago [9–14]. These studies also made converging observations regarding the potential neural loci for integration of optic flow, including the interpeduncular nucleus, medulla oblongata, and inferior olive.

In contrast, it remains disputed why the fish integrates optic flow for such a long time. Some argued that this integrative process allows fish to estimate the direction of optic flow more accurately in the presence of noise (“evidence accumulation”) [9, 11]. Others claimed that fish are keeping track of their position in the environment by integrating optic flow over time (“position homeostasis”) [14]. Although the two functional interpretations make different predictions as to how self-generated optic flow should affect the OMR, these have not yet been tested experimentally.

The primary goal of the present study is to dissect the algorithms involved in the second-scale integration of optic flow to better understand their functions. First, by manipulating the gains of visual feedback, we demonstrate that the OMR depends only on the history of the externally generated (i.e., exafferent) optic flow, ignoring the self-generated (i.e., reafferent) optic flow. This pattern holds for both position- and heading-stabilizing OMR and is more consistent with the evidence accumulation interpretation. Second, we characterize the dynamics of the history dependence in the OMR using reverse correlation and delay-based paradigms. Unexpectedly, the two approaches uncovered different timescales, suggesting that the fish becomes more forgetful in a more dynamic environment. With whole-brain calcium imaging with light-sheet microscopy, we demonstrate that the optic flow selective cells across the brain receive motor efference copy signals, such that they respond less to the reafferent optic flow. Finally, with two-photon calcium imaging, we show that neurons in the inferior olive integrate forward and backward optic flow separately with long time constants, providing clues to how the multiple timescales of the history dependent OMR is implemented.

## Results

### Forward OMR is dependent on sensory history

To probe the optomotor response (OMR) and its history dependence in the zebrafish larvae, we utilized a head-restrained, closed-loop behavioral assay as described before [15] (see Materials and Methods for details). Briefly, 6 to 8-day-old larvae were head fixed in low-melting point agarose with tails free to move and observed visual stimuli projected below with a small projector (Figure 1A). The movements of the tail were monitored with a high-speed camera, which was then used to infer intended swim patterns and to provide real-time visual feedback. In the first set of experiments, we focused on forward swimming in response to forward optic flow. Visual patterns flowing forward give the impression that the observer is drifting backward relative to the environment and elicit robust OMR in larval zebrafish [15]. Throughout the paper, we will describe optic flow in terms of the directions of movements of the visual patterns seen from the fish, as opposed to the directions of the implied observer movements.

**Figure 1:**
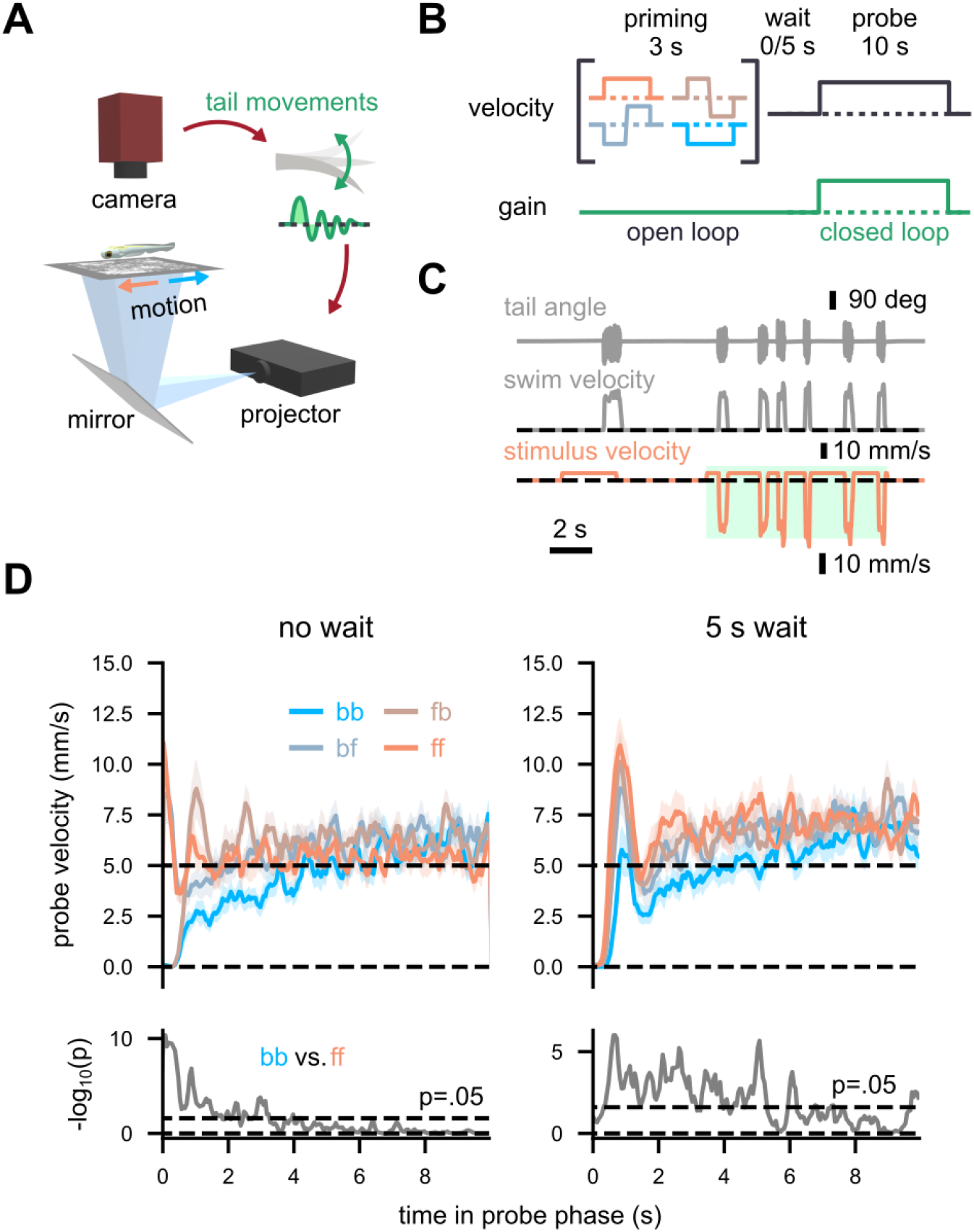
Forward optomotor response depends on the history of optic flow. (A) A schematic of the behavioral setup. (B) A schematic of the priming-delay assay. (C) An example trial from a single fish. The dotted horizontal lines indicate 0. The green-shaded region represents the closed-loop probe phase. (D) (top) Average swim speed in the probe phase color-coded by priming conditions, and (bottom) −log10 (p values) between forward and backward priming conditions from time point-wise paired t-test. Trials with and without the 5 s delay period are plotted separately. The solid lines and the shaded areas respectively indicate the mean across fish and its standard error. N = 30 fish.

In the initial experiment, we performed an optic flow memory assay similar to the ones used in [14] (Figure 1B). In each trial, fish first observed three seconds of visual flow moving either forward or backward at 5 mm/s, with a possible directional switch in the middle (“priming” phase). This priming phase was in an open loop: that is, tail movements did not affect the ongoing visual stimuli. After an open-loop delay period without visual motion for either 0 or 5 seconds, the fish then entered the probe phase, where forward flow at 5 mm/s was presented for 10 seconds. During the probe phase, the tail beats of the larvae were translated to backward movements in the visual patterns (i.e., closed-loop) (Figure 1C) (see Methods for the details). We observed statistically significant effects of the priming directions on the swimming velocities during the probe phase (Figure 1D), revealing the history dependence of the forward OMR. In particular, the instantaneous swim velocity was higher in the forward priming conditions compared to the backward condition up until several seconds into the probe phase, for both 0 and 5 s waiting conditions (Figure 1D). The direction-switching priming conditions resulted in intermediate behaviors. These results essentially replicate the observations by [14], but in non-paralyzed fish.

### Self-generated optic flow does not count toward the future OMR

Optic flow can arise as a consequence of voluntary movements of the observer (i.e., reafference) or from involuntary movements due to environmental perturbations (i.e., exafference), such as water flow or wind. If the purpose of the integration of optic flow is to accurately estimate the velocity of the environmental flow (“evidence accumulation”) [9, 11], then only exafferent, but not reafferent optic flow should be integrated. Alternatively, if fish are keeping track of their position relative to the environment by integrating optic flow (“position homeostasis”) [14], then they should integrate both reafferent and exafferent optic flow. To disambiguate these two functional interpretations, we next explicitly manipulated the reafferent optic flow while keeping the exafference constant (Figure 2A). Each trial started with 5 seconds of the “priming” phase, where 5 mm/s forward optic flow was presented. After a delay of 2 seconds, 5 mm/s forward optic flow was presented again for 10 seconds (“probe”). Here, the priming phase was in a closed loop and the probe phase was in an open loop, unlike in the previous experiment (Figure 1). In addition, the gain of the closed-loop feedback was either increased to 1.5 or decreased to 0.5 in some trials. Thus, the fish always received a fixed amount of exafferent optic flow in each priming phase, but different amounts of reafferent optic flow, depending on their behavior and the closed-loop gain. We made sure that the fish were aware of this gain manipulation by comparing the bout durations (Figure S1).

**Figure 2:**
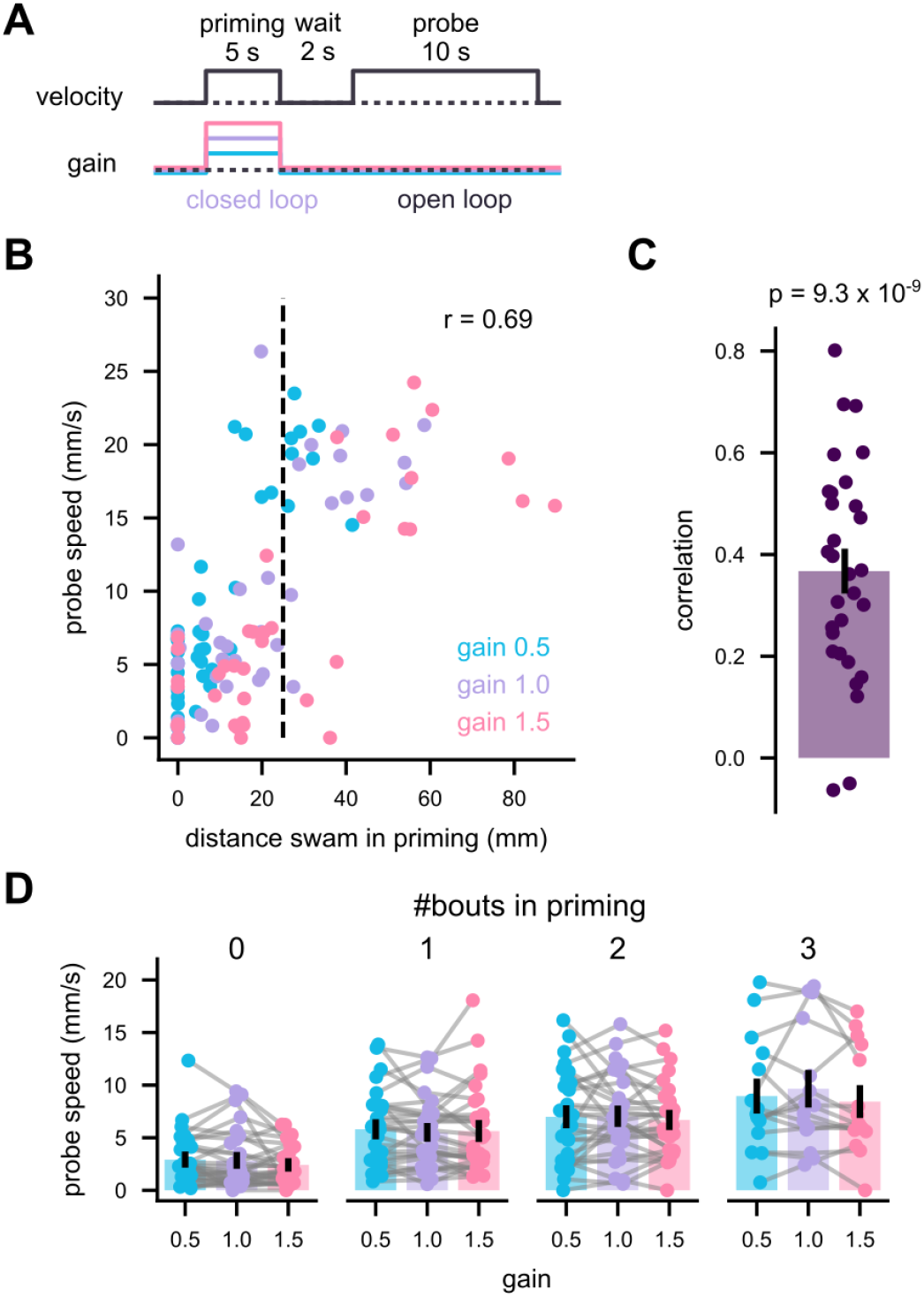
Forward optomotor response depends on the history of exafferent optic flow. (A) A schematic of the protocol. (B) Data from an example fish. Average swim speeds during the probe period (assuming 1.0 gain) are plotted against the distance swam during the priming period. Each dot corresponds to a trial. (C) Pearson correlation between the probe speed and priming swim distance from all fish. N = 30 fish. (D) Trial-averaged probe swimming speed as functions of the priming gain and the number of bouts during the priming. The bars indicate the average across fish, and the dots individual fish. The data from identical fish are connected. For a given number of priming bouts, no pair of gain combinations exhibited significantly different probe speeds (p > 0.05). N = 30, 30, 29, 15 for # bouts = 0, 1, 2, 3. N is smaller than 30 in some conditions because some fish did not have trials with the same number of bouts across all three gain conditions. P-values are from signed-rank tests.

Figure 2B shows the data from a representative single fish. In this fish, more swimming in the priming phase led to more swimming in the probe phase. This positive correlation between the priming and probe phase was consistent across the tested population (Figure 2C). Note that neither of the aforementioned functional hypotheses predicts the positive priming-probe correlation: the evidence accumulation hypothesis predicts that the reafferent optic flow seen during the priming phase would be ignored for the future OMR (i.e., no correlation). The position homeostasis hypothesis predicts that more swimming in the priming phase would lead to less swimming in the probe phase (i.e., negative correlation).

A plausible source of this positive correlation is a slow fluctuation in the alertness of the fish. For example, in some trials, fish might be more alert in general and swim more in both phases and vice versa. To isolate the effect of the reafferent optic flow from the effect of alertness, we sorted the trials by the number of swim bouts in the priming phase (Figure 2D). Even among the trials with the same number of priming bouts, we still did not observe the effects of the priming phase closed-loop gain on the probe phase swimming. In other words, the history-dependent OMR does not depend on the reafferent optic flow. This observation is more consistent with the evidence accumulation hypothesis, rather than the position homeostasis.

### The rotational OMR also depends only on exafferent optic flow

The experiments so far established that the forward OMR is dependent on the history of exafferent, but not reafferent optic flow. We next wondered whether the same holds for other types of OMR. Rotational optic flow in yaw directions triggers whole-body rotation in various teleost fish [16, 17], which we will refer to as the rotational OMR. First, we established that the head-restrained zebrafish larvae show robust rotational OMR, whose magnitude was roughly proportional to the stimulus angular velocity, albeit with a low gain (Figure S2A-C). Next, we performed a priming-delay assay similar to Figure 1B in the rotational dimension (Figure S2D). We found that when the priming stimulus rotated in the same direction as the probe stimulus (i.e., ipsi-directional), the fish performed rotational OMR more, compared to when the probe moved in the opposite direction (i.e., contra-directional) (Figure S2E). This effect was significant even after waiting for 5 seconds, establishing the history dependence of the rotational OMR.

We then asked whether the fish integrated only exafferent or total rotational optic flow for the rotational OMR with a gain-manipulation assay similar to Figure 2. This was of particular interest because the brain of larval zebrafish is equipped with head direction neurons [18], which is in principle suited for implementing a homeostatic control of the head direction. Each trial started with a 5-second-long closed-loop priming phase, where 30 °/s rotational optic flow in either direction was presented. Followed by a 2-second waiting period, another 10 seconds of 30 °/s rotation was presented in an open-loop (“probe”) (Figure 3A). The gain of closed-loop feedback in the priming phase was either low (0.5) or high (1.5). Consistent with the prior experiment (Figure S2D, E), the ipsi-directional priming resulted in more rotational OMR in the probe phase (Figure 3B), while the effect of feedback gain was qualitatively not apparent. We then analyzed how behaviors during the priming phase affected swimming in the probe phase on a trial-by-trial basis, focusing on the ipsi-directional priming conditions. Figure 3C shows example data from a single fish, similar to Figure 2B. In this fish, more ipsi-directional turning in the priming phase led to more rotational OMR in the probe phase. This positive correlation was statistically significant across the tested population. This effect is again not predicted by the evidence accumulation or position homeostasis hypotheses and can be parsimoniously explained by fluctuations in the alertness of the animals. To isolate the effect of the gains, we stratified trials by cumulative turn angles in the priming phase into quartiles, disregarding gains. In each quartile, fish performed similar amounts of lateralized tail bends, but received reafference three-fold different depending on gains. Yet, in any quartile, increased reafference did not result in reduced probe OMR predicted by the homeostatic hypothesis. The results from the contra-directional priming conditions are more complex due to the fish’s tendency to turn consecutively in the same direction11, but similarly, we did not detect any effect of the gains (Figure S3). Overall, the rotational OMR, similar to the translational OMR, appears to depend on the history of exafferent optic flow.

**Figure 3:**
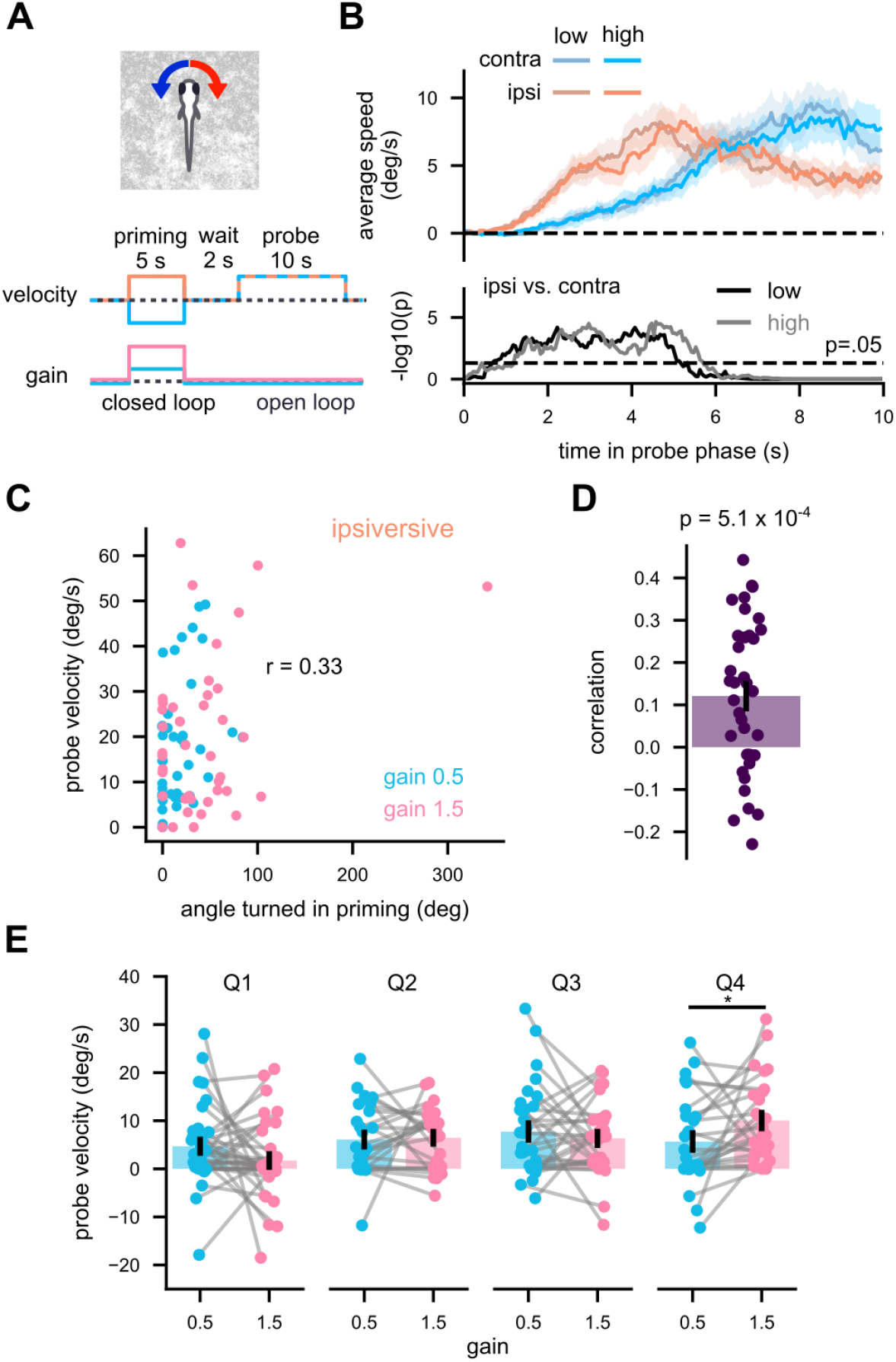
Rotational OMR depends on the history of exafferent optic flow. (A) A schematic of the protocol. (B) (top) Average turning velocity during the probe phase and (bottom) p-values between ipsi- and contra-directional conditions within each gain condition over time. The horizontal dotted line corresponds to p = 0.05. The solid lines are the averages across fish, and the shaded areas indicate the standard errors of the means. (C) Data from an example fish in ipsi-directional priming conditions. Turn velocities during the probe period (assuming 1.0 gain) are plotted against the angle turned during the priming period. Each dot corresponds to a trial. (D) Pearson correlation between the probe speed and priming swim distance from all fish. The bar indicates the average across fish, and the dots individual fish. N = 36 fish. (E) Trial-averaged probe turning velocities stratified into quartiles based on the cumulative tail angles during the priming phase. The data from identical fish are connected. Only data from ipsi-directional conditions are plotted. See Figure S3 for the contra-directional data. Only the fourth quartile (Q4) had a significant effect of the gain, but in the opposite direction as expected from the homeostatic hypothesis. N = 35, 25, 29, 27 for Q1, 2, 3, 4. The p-values are from (B) time point-wise paired t-tests or (D, E) signed-rank tests.

### Reverse correlation reveals optic flow memory lasting for a couple seconds

The experiments so far established that both forward and rotational OMR depend on the history of exafferent optic flow. Next, we aimed to characterize the kinematics of the history-dependent OMR in an unbiased manner using a reverse correlation technique [19, 20]. We presented head-embedded fish with a visual pattern moving either forward or backward at 15 mm/s in an open-loop fashion. The direction of the motion flipped stochastically following a Poisson point process with the rate of 0.5 Hz (Figure 4A). We then calculated the average of stimulus velocity time traces leading up to forward swimming bouts, weighted by the peak speed of the bouts (see Materials and Methods for details). We refer to this weighted average trace of velocity as a bout-triggered average (BTA) stimulus, which can be interpreted as an estimate of the filtering processes happening in the larval brain.

**Figure 4:**
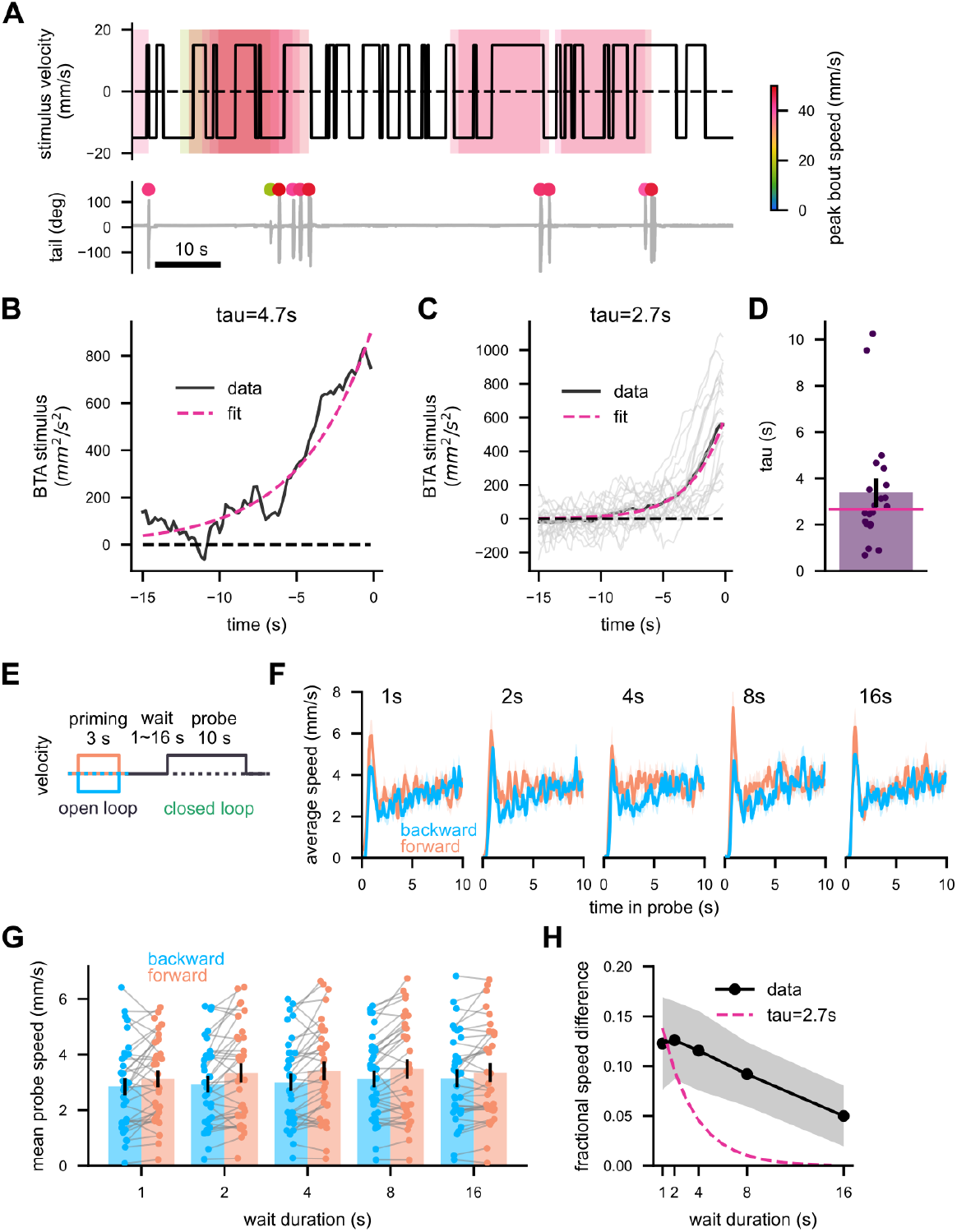
Multiple timescales of history-dependent OMR. Example traces of (top) stimulus velocity and (bottom) tail angles during the reverse correlation experiment. Bouts and velocity traces leading up to them are indicated by dots and boxes colored according to the bout speed. (B) The bout triggered average (BTA) velocity trace from the fish in (A) (black), and an exponential fit (magenta). (C) The BTA stimuli from individual fish (gray) as well as their population average (black). A decaying exponential was fit to the group average (magenta), yielding the time constant of 2.7 s. (D) The time constants of the exponential fits to the individual BTA (purple), as well as the time constant of the exponential fit to the population average (magenta line). (E) A schematic of the long delay experiment. (F, G) The population-average swim velocity in the probe phase (F) over time and (G) time-averaged, color-coded by the direction of the priming stimuli and sorted by waiting durations. (H) Fractional differences in the probe swim speed by the conditioning directions, as a function of the wait duration (black), alongside an exponential with the 2.7 s decay constant (magenta). (C, D) N = 21 fish. (F-H) N = 35 fish. Each dot in scatter plots indicates a single fish, the bars indicate averages across fish, and the error bars are standard errors of the mean. The solid line and the shaded areas indicate the mean across fish and its standard error.

Figure 4B, C shows BTA stimuli from an individual fish as well as averaged across the population. The population average BTA was fit well by a decaying exponential with the time constant of 2.7 seconds (Figure 4D). We performed an equivalent experiment for the rotational OMR, and obtained a comparable time constant (Figure S4A-D). We made sure with a simulation that this time constant does not simply result from the autocorrelation of the stimulus (Figure S4E). Overall, it appears that both forward and rotational OMR have history-dependent components with the time constant of around 3 seconds.

### The conditional leak model captures the two timescales of optic flow integration

While the time constant of around 3 seconds is in line with some previous reports [9, 11, 12], others have observed much longer history dependence spanning tens of seconds [14]. To make sure that this large discrepancy in timescales did not result from differences in experimental setups, we repeated the priming-delay assay as in Figure 1 with various delay durations between 1 to 16 seconds (Figure 4E, F). We then plotted the difference in the probe swim speed between forward- and backward-priming conditions as a function of the delay durations (Figure 4G, H). We found that the probe speed difference by the priming direction only fell from 12% to 6% after 16 seconds of delay, a kinematics much slower than the reverse correlation-based BTA.

What difference between the two experiments (Figure 4A, E) led to these dramatic differences in the observed timescales of the OMR? In the reverse correlation experiment, the fish continuously observed optic flow changing directions, while in the priming-delay assay, the fish did not see any visual motion in the delay phase. Thus, we hypothesized that the presence of concurrent optic flow makes the fish more forgetful about the past optic flow. To express this hypothesis more quantitatively, we built a simple model which we refer to as the conditional leak model (Figure 5A) (See Materials and Methods). Similar to the previous models [9, 11] our model unit receives the velocity of optic flow as its input, and integrates it in a leaky fashion. The activity of the model unit is subjugated to a static, bounded nonlinearity to generate the rate of stochastically generated swim bouts. In addition, the leakiness is increased if the input speed is above a certain threshold, allowing the system to have different time constants in the presence and absence of optic flow. By appropriately choosing the model parameters, the conditional leak model can replicate a reverse correlation-based BTA with a relatively short time scale (Figure 5B, C) as well as a longer history dependence in a simulated priming-delay assay (Figure 5D, E), as expected.

**Figure 5:**
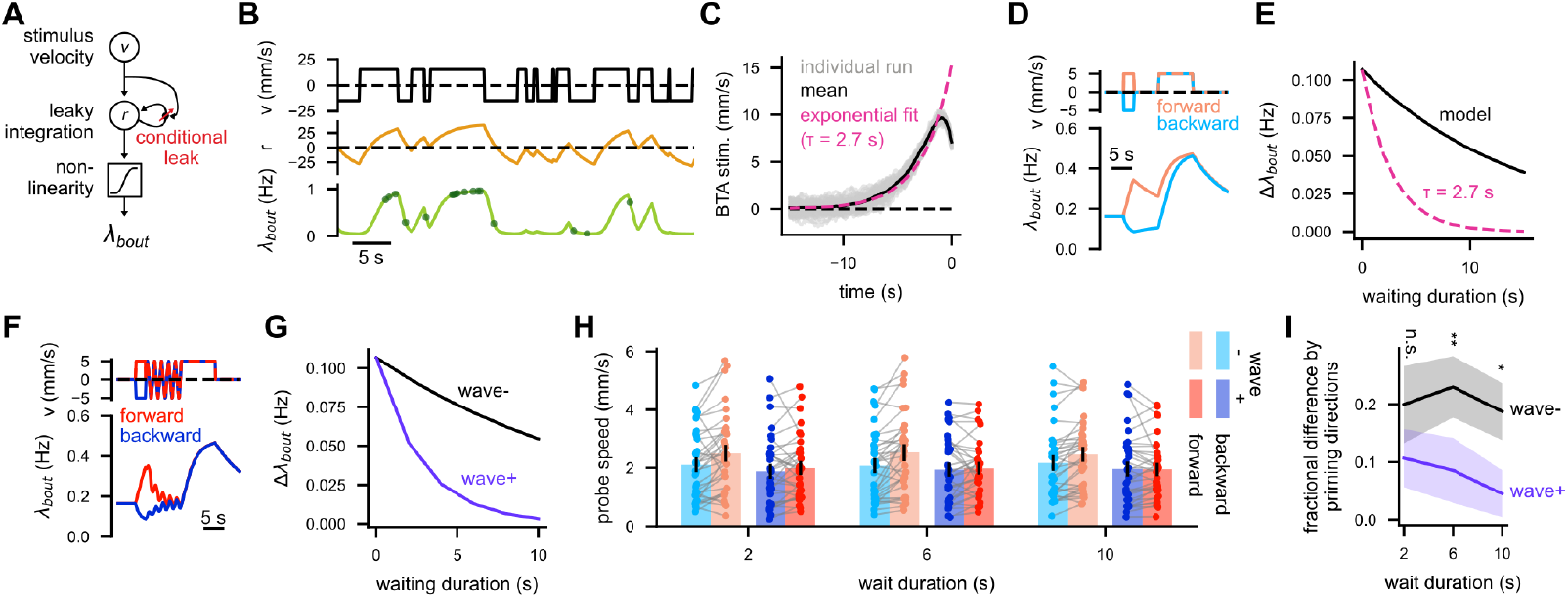
The conditional leak model reconciles the two timescales of OMR. (A) A schematic of the conditional leak model. (B) An example response of the model to the stochastic direction flip stimulus as in Figure 4A. The yellow line represents the model neuron response, and the green line the bout rate after the nonlinearity. The green dots represent simulated bouts. (C) Reverse correlation-based BTA stimulus from the model. (D) Bout rate λ_bout_ calculated by the model in response to the priming protocol. (E) The difference in mean λ_bout_ during the probe phase by the priming direction as a function of waiting time, as in Figure 4H. (F, G) Same as (D, E), but with the “wave” with net-zero displacement added to the waiting phase. The wave opens the “gate” in the model and reduces the effective time constant of the integrator. (H) Time-averaged swim speed of real fish during the probe phase as a function of waiting duration, sorted by priming durations and wave conditions. (I) Fractional differences of the probe swim speed by the priming directions, as functions of waiting durations, sorted by the wave conditions. Each dot in scatter plots indicates a single fish, the bars indicate averages across fish, and the error bars are standard errors of the mean. The solid line and the shaded areas indicate the mean across fish and its standard error. N = 34 fish, n.s.: not significant (p > .05), *: p < .05; **: p < .01. P-values are from signed-rank tests.

A non-trivial, testable prediction from the conditional leak model is that even a sequence of optic flow that imply net-zero displacement can affect the future OMR by making the fish more forgetful. For example, if we inject a sinusoidally oscillating pattern of velocity inputs with the zero mean to the conditional leak model during the delay phase of the priming-delay assay (Figure 5F), the probe bout rate difference by priming directions decay much faster, following the shorter time constant (Figure 5G). In the real fish, we also observed that oscillating the visual pattern back and forth during the delay phase makes the fish more forgetful (Figure 5H, I). For example, the forward OMR of fish was barely affected by the priming directions when the delay phase was longer than 6 seconds. In a separate experiment, we tested how various types of dynamic visual patterns affect the decay of the optic flow memory (Figure S5). In summary, we found that the optic flow integration in the larval zebrafish involves multiple timescales controlled by the existence and absence of concurrent optic flow.

### Direction selective cells across the brain receive efference copy signals

One of the key insights from the behavioral experiments above is that the fish OMR depends on the history of the exafferent optic flow [2] (Figure 2, Figure 3). This implies that the responses of optic flow-tuned neurons to expected reafference are canceled by motor efference copy signals, somewhere along the pathway generating OMR. To identify such efference copy signals in the brain, we performed a brain-wide calcium imaging experiment.

Here, we imaged the brains of the larvae broadly expressing a calcium indicator in a nuclear-localized fashion (elavl3:H2B-GCaMP6s) with a light-sheet microscope (Figure 6A). At the beginning of each experiment, we presented the fish with visual patterns moving in the four directions (forward, backward, left, right) repeatedly, and selected neurons that reliably responded to them in a direction selective fashion (see Methods for the detailed selection criteria). This revealed stereotyped clusters of direction selective neurons in the pretectum, anterior hindbrain, and inferior olive, as have been previously observed (Figure 6B) [21–23]. During the rest of the experiment, the fish experienced a slow (5 mm/s), continuous forward optic flow. Every time the fish made a swim bout, the fish stochastically received visual feedback (i.e., backward flow) or no feedback. The dynamics and peak velocity of the feedback flow was fixed, regardless of the properties of the swim bout (see Methods). In addition, backward flow with the same dynamics as the fixed feedback was sometimes presented in open-loop, in the absence of swims. We then calculated the change of fluorescence for each cell after the three types of events (i.e., swims with or without feedback and backward flow without swims) (Figure 6C).

**Figure 6:**
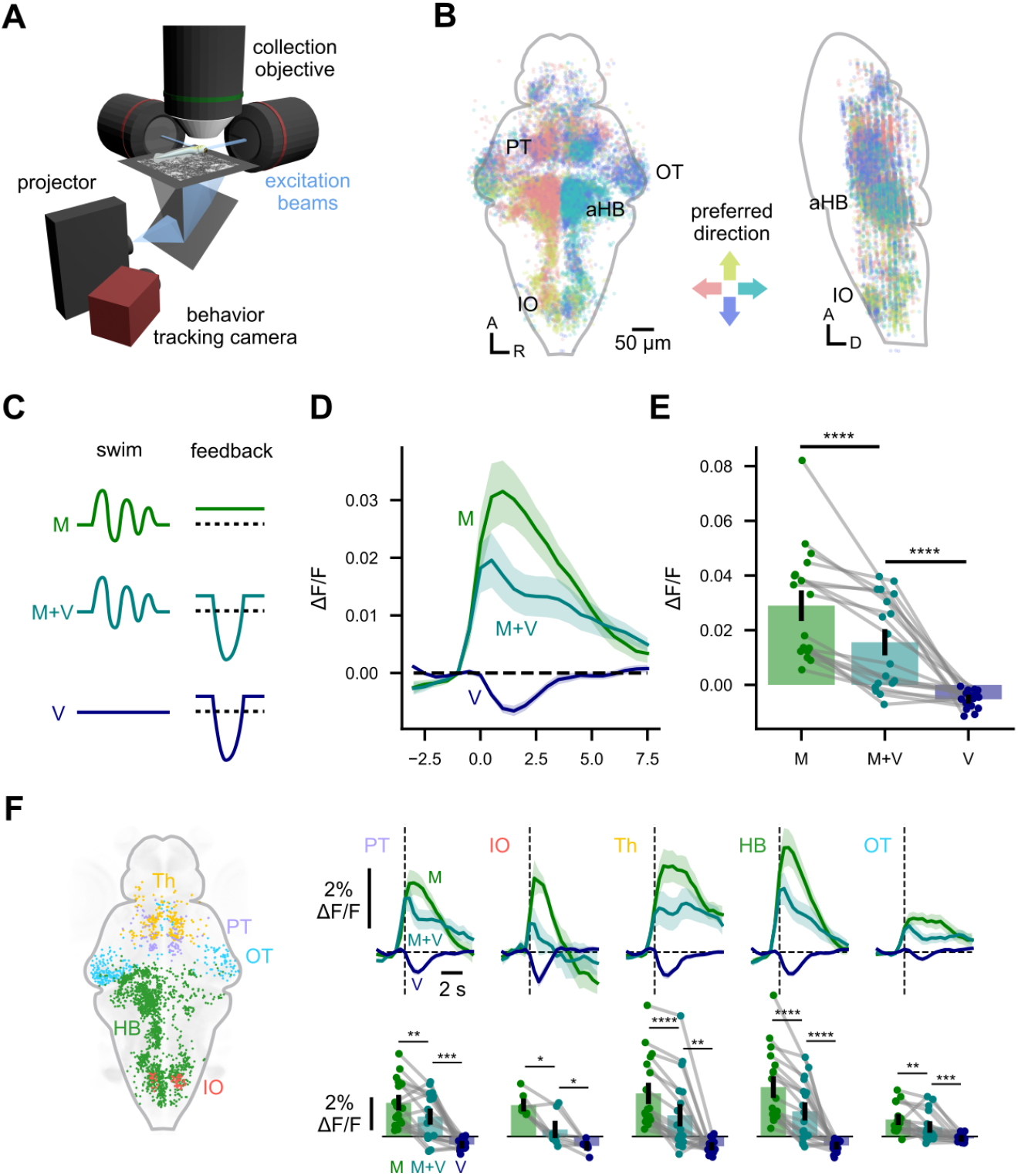
Neurons tuned to forward optic flow receive positive efference copy signal. (A) Schematics of the light sheet microscope. A pair of excitation objectives delivered perpendicular sheets of excitation beam, which moved in the Z direction in synchrony with the collection objective, allowing fast volumetric imaging. (B) Direction selective neurons identified by grating stimuli in the (left) horizontal and (right) sagittal views. Pooled across N = 18 fish. (C) Schematics of the experiment. Under the presence of a constant forward optic flow, fish swam in an open-loop (M), swam and received a fixed backward feedback (M+V), or received the backward flow in an open-loop (V). (D) Calcium activity triggered by the three types of events, averaged across the all forward-tuned cells. The solid lines and shaded areas respectively indicate the mean across fish and its standard error. (E) Same as (D), but averaged within the 3 second window after the triggering event. The bars indicate the averages across fish. Each dot represents a single fish, and the data from the same fish are connected by lines. (F) (top) Same as (D), but restricted to cells in specific brain regions as color-coded in the left image. (bottom) Same as (E), but for each brain region. N = 17, 7, 18, 18, 18 fish for PT, IO, Th, HB, OT. *: p < .05, **: p < .01; ***: p < .001; ****: p < .0001 from signed-rank tests. A: anterior; R: right; D: dorsal; PT: pretectum; IO: inferior olive; Th: Thalamus; HB: hindbrain; aHB: anterior hindbrain; OT: optic tectum.

Figure 6D shows the triggered activity of forward selective neurons averaged across the entire brain. Backward flow without swims resulted in a transient dip in the activity of forward selective cells. Interestingly, swims without feedback resulted in positive activity. This observation is not trivial, given that these cells are selected purely based on their visual response properties. The motor-driven positive activities and the dip in activity due to backward flow worked to cancel each other after the swims with feedback. Consistent patterns were observed when we restricted our analyses to specific brain regions, including those that have been implicated in OMR generation, such as pretectum and inferior olive (Figure 6E). We also found backward-tuned cells whose activity is suppressed after swim bouts (Figure S6), suggesting directional specificity of the efference copy signals. In summary, optic flow selective neurons supposedly serving the forward OMR receive the motor efference copy such that they respond less to reafference, mirroring the behavioral observation that forward OMR depends on the history of the exafferent optic flow.

### Oscillating stimuli induce additive baseline shifting in the olivary memory cells

In the final experiment, we attempted to gain insights into how multiple timescales of the optic flow memory are implemented in the brain. To do so, we decided to focus on the inferior olive (IO). A previous study has shown that the IO neurons receive information about the integrated optic flow from the medulla oblongata, and that they are necessary for the memory-based forward OMR [14]. Here, we recorded activities of IO neurons expressing GCaMP6s under the control of vglut2a:Gal4 [24] using a two-photon microscope (Figure 7A). In each trial, fish observed a visual pattern moving forward or backward for 2 seconds at 5 mm/s (i.e., “priming” pulse) (Figure 7B). In a subset of the trials, the priming pulse was followed by 6 seconds of sinusoidal, back-and-forth oscillations of the pattern (peak 5 mm/s, 0.5 Hz), as in Figure 5F.

**Figure 7:**
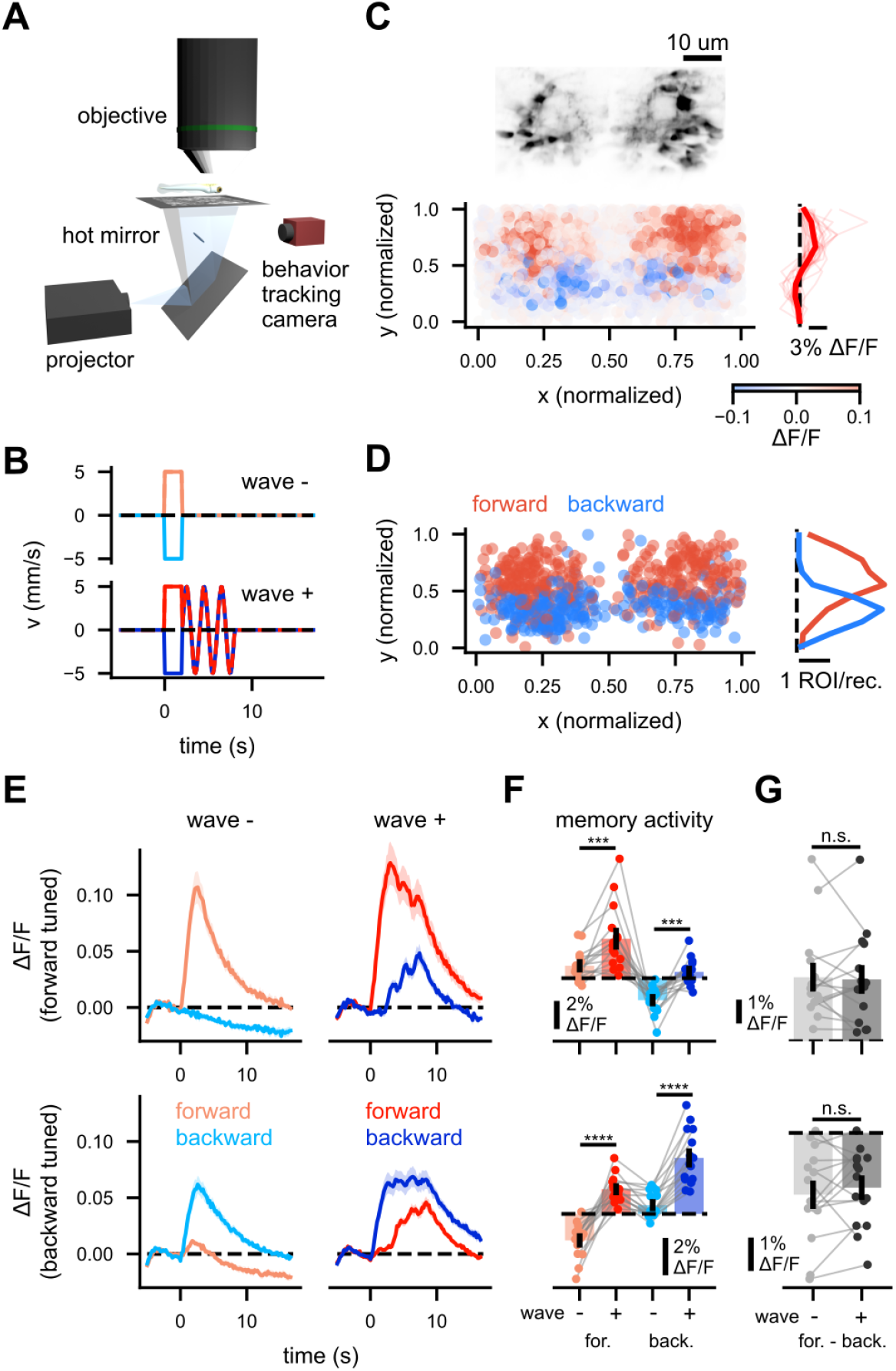
Increased baseline activities of the olivary memory cells in a dynamic visual environment. (A) Schematics of the two photon microscope setup. (B) Schematics of the visual stimuli. The fish received 2 seconds-long priming pulse in forward or backward directions, optionally followed by sinusoidal oscillations of the visual pattern (i.e., “wave”). (C) (top) Example image of the inferior olive (IO) labeled by vglut2:Gal4. (bottom left) The IO ROIs color-coded by the difference of their response to forward and backward priming pulses. Pooled across 26 recordings in 15 fish. (bottom right) The median ΔF/F during the priming pulse, as a function of normalized Y (i.e., anterior-posterior) position. The bold and thin lines respectively indicate the average across 26 recordings and individual recordings. (D) (left) The normalized positions of the ROIs that had reliable memory activity, color-coded by their preferred priming pulse directions (see Methods for the detailed selection criteria). (right) The histogram of the memory cells by the normalized Y position, separately for forward and backward tuned cells. (E) The response time traces of the (top) forward- and (bottom) backward-tuned memory cells to the stimuli in (b). Notice how the oscillating stimuli increase the activity of the both functional cell types. The solid line and shaded areas represent the mean across fish (N = 15) and its standard error. (F) The memory activity (i.e., the mean ΔF/F between 8 to 17 s from the priming pulse onset) by stimulus type, for the (top) forward and (bottom) backward selective memory cells. Each dot represents an individual fish, and the data from the same fish is connected by grey lines. (G) The difference of the memory activity by the priming pulse direction, with or without the oscillating stimuli, for the (top) forward and (bottom) backward selective memory cells. n.s.: not significant; *: p < .05, **: p < .01; ***: p < .001; ****: p < .0001 from signed-rank tests.

First, we observed that the IO cells that positively responded to the forward and backward priming pulses respectively tended to be positioned anteriorly and posteriorly, replicating the recently reported topographic pattern [25] (Figure 7C). Next, using a half of the data from the trials without the oscillations, we identified IO cells whose activities reliably reflected the priming pulse direction more than 6 seconds after its offset (hence-forth “memory cells”; see Methods for details). The distribution of the forward- and backward-tuned memory cells exhibited a topographic pattern congruent to those during the priming pulse (Figure 7D). We then examined the dynamics of the responses of the memory cells using the remaining portion of the data not used for selecting those cells. Contrary to the divisive behavior of the conditional leak model (Figure 5F), the oscillatory visual motion additively increased the activity of the IO memory cells regardless of their directional preference (Figure 7E, F). The increase in the forward and backward memory cell activities appeared mostly parallel (Figure 7G), and thus the trace of the priming pulse direction is clearly discernible from the memory cell activities even after the oscillatory inputs. However, these parallel shifts could lead to earlier forgetting depending on the nonlinearity of the downstream circuitry. Example quantitative implementations of this mechanisms are provided in Figure S7. Overall, the observation here suggests that additive baseline shifting and nonlinearities downstream of the IO implement the shortening of the OMR hysteresis in a dynamic visual environment.

## Discussion

The primary goal of the present study was to gain a better insight into the algorithms underlying the seconds-scale history dependence in the larval zebrafish OMR. Through a series of behavioral experiments, we demonstrated that (1) the OMR depends on the recent history of exafferent, as opposed to net, optic flow, and that (2) the timescale of the OMR hysteresis is shortened in a more dynamic visual environment. Paralleling these behavioral observations, we observed that (1) the neurons directionally tuned to the optic flow throughout the brain receive motor-related signals that tend to cancel the responses to reafference, and that (2) the optic flow memory in the IO is attenuated by additive baseline shifts and nonlinearity in the face of dynamically changing optic flows.

As discussed in the introduction, previous studies have postulated different functional goals for the hysteresis in the zebrafish OMR: Some argued that the fish integrate optic flow to better estimate the velocity of the environmental flow in the face of noisy visual inputs (“evidence accumulation”) [9, 11], while others suggested that fish perform proportional-derivative control over their position using the visual motion signals (“position homeostasis”) [14]. The observation that the reafferent visual motion does not count towards the future OMR drive (Figure 2, Figure 3) favors the evidence accumulation interpretation, because estimating the position requires integrating the visual motion regardless of its sources. In light of these findings, it seems unlikely that the medullar and olivary neurons causally connected to the history dependent OMR [14] are truly encoding the animals’ position similar to the head direction [18] and place cells [26] found elsewhere in the larval zebrafish brain.

How is the cancellation of the expected reafferent optic flow implemented? In the whole brain imaging experiment, we observed efference copy-like signals in diverse brain regions selective for optic flow directions (Figure 6). A parsimonious hypothesis is that the subtraction of the expected reafference happens in relatively peripheral optic flow detectors (e.g., in the pretectum), which then propagate the computed exafferent signal to the other regions. Such arrangement has been found in the optomotor circuitry in Drosophila [27] and the avian pretectum [28]. Alternatively, multiple optic flow sensitive brain regions in the brain might receive efference copy signals in parallel, according to their individual functional needs. Another open question is the source of the efference copy signals. A prime candidate is the cerebellum, which has been considered to implement internal models to predict sensory outcomes of the motor commands (i.e. forward models) [29]. In fact, in the specific context of the larval zebrafish OMR, we have previously shown that the cerebellar Purkinje cells are required for the fish to learn a new association between their swim bouts and the resultant visual feedback [12]. In addition, a recent study suggested that the inputs from the torus longitudinalis cancel self-generated visual inputs in the optic tectum [30], which might explain the negative bout-triggered activities in the tectal neurons (Figure S6).

Another major behavioral finding of the present study is that the timescale of the optic flow memory depends on the stimulus dynamics (Figure 4, Figure 5). Although the behavioral results do not distinguish whether the shortening of the memory timescale is caused by the existence of optic flow itself or the changes of its directions, the mechanisms suggested by the IO physiology (Figure 7, Figure S7) favors the second possibility. Viewed through the lens of the evidence accumulation, it makes intuitive sense to be more forgetful in a rapidly changing environment, as past sensory evidence would become more quickly irrelevant to the current state which the animal is trying to estimate. Indeed, studies on normative models of decision making have shown that ideal observers discard older evidence in a manner dependent on the environmental volatility [31]. Similarly, temporal filtering in the retinal ganglion cells become faster in the face of stimuli with higher variance [32].

In our recordings from the IO neurons, we observed that visual patterns oscillating back-and-forth increase the activity of both forward- and backward-tuned memory cells. The increase in the average activity in the face of stimuli with higher variance naturally results from the nonlinearity of neurons (as modeled in Figure S7C), and have been widely observed in early visual neurons in both vertebrates [32] and insects [33]. Yet, this is somewhat surprising in the context of the optic flow detection. Optic flow tuned neurons across taxa (e.g., primate MST [34], mammalian [35] and avian [36] pretectum, lobula plate tangential cells in flies [37]) are typically inhibited by the optic flow opposite to what they prefer (i.e., motion opponency), thereby representing self-motion in a signed, approximately linear fashion. The algorithmic models of the zebrafish history dependent OMR, including our initial model (Figure 5), also implicitly or explicitly assume motion opponency before the integration step [11, 12, 14]. Our tentative functional interpretation of this unexpected lack of motion opponency in the IO memory cells is that, by keeping track of forward and backward motion separately, they allow the downstream circuitry to easily construct an estimate of the environmental volatility (as in Figure S7E). It is of future interest to examine how the downstream circuitry (i.e., cerebellar Purkinje cells) transform these nonlinear memory signals in the IO.

## Acknowledgments

RT was supported by the European Molecular Biology Organization (EMBO ALTF 732-2022) as well as the Human Frontier Science Program (HFSP LT0027/2023-L) for this work. This research was funded by the German Research Foundation (DFG) under Germany’s Excellence Strategy within the framework of the Munich Cluster for Systems Neurology (EXC 2145 SyNergy, identifier 390857198), through the “Enhanced resolution microscopy” project DFG – Projektnummer 518284373, by the Volkswagen Stiftung via a Life? grant, and by the Max Planck Gesellschaft.

## Contributions

RT and RP conceived the project. RT performed the experiments, analyzed the data, and performed the simulations. RT and RP wrote the manuscript.

## Competing interests

The authors declare no competing interest.

## Materials and Methods

### Zebrafish Husbandry

All experimental procedures were conducted according to protocols by the Technische Universität München (TUM) and the Regierung von Oberbayern (animal protocol number 55-2-1-54-2532-10112 and 55.2-2532.Vet_02-24-5). Adult zebrafish (Danio rerio) were housed in the fish facility in the Institute for Neuronal Cell Biology at TUM. The adult fish were maintained in water temperature of 27.5 - 28.0 °C on the 14:10 hour light:dark cycle. All experiments were performed on 6 to 8 days post fertilization (d. p. f.) larvae of undetermined sex. The Tüpfel Long-fin (TL) strain was used for all behavioral experiments (Figures 1-5, S1-5). For the whole-brain imaging experiment (Figure 6), Tg(elavl3:H2B-GCaMP6s)jf5 was used [38]. For the IO imaging experiment (Figure 7), fish carrying Tg(vglut2a:Gal4)nns20 [24] and Tg(UAS:GCaMP6s)mpn101 [39] were used. Fish with mutant background (mitfa-/-) lacking melanophores were used for the imaging experiments to allow optical access to the brain.

### Behavioral Experiments

For head-fixed behavioral experiments, larvae were embedded in 1.5% low melting point agarose (Thermo Fisher) in 35 mm Petri dishes. Fish were immersed in water from the fish facility, and agarose around the tail was carefully removed so that the larvae can beat their tails. Experiments were performed at least one hour after the embedding procedure to allow fish to acclimate to the head-restrained condition. The movements of the tail were monitored with high-speed cameras at above 200 Hz under infra-red illumination. Visual stimuli were projected below the fish with compact LCD projectors at approximately 60 Hz. The viewing distance was about 5 mm. Real-time tracking of the tail movements and visual stimulus generation was performed with the stytra package [40].

To infer the intended swimming patterns of head-restrained fish, movements of the tail were analyzed as follows [40]: In each video frame, the tail of a larva was first segmented into 7 to 10 linear segments. Next, the sum of angular differences between all neighboring segment pairs were calculated, which added up to the angle between the body and the tip of the tail. We then calculated the instantaneous swim vigor of the fish as the standard deviation of this tail angle within a 50 ms rolling window. A swim bout is defined as an episode of vigor above 0.05 rad. As a proxy of turning angle at each bout, we calculated the bout bias as the cumulative sum of the tail angle within the first 70 ms in the bout, subtracted with the baseline tail angle 50 ms before the bout onset [41].

To simulate translational and rotational optic flow, self-repeating spatial pink-noise patterns [18] were rigidly translated or rotated about the fish. Exact protocol structures are compiled in Table S1. In the closed-loop experiments, the translational and angular velocities of the stimuli were manipulated as follows: In forward OMR experiments (Figure 1, Figure 2, Figure 4, Figure 5, Figure S5), forward swimming velocity of the fish was estimated by multiplying the basal gain of 30 mm/s/rad on the vigor. In the closed-loop epochs, this swim velocity was subtracted from forward stimulus velocity during swim bouts. Additional gains of 0.5 or 1.5 were applied on the forward velocity in some experiments (Figure 3, Figure S2). In the closed-loop epochs of the rotational OMR experiments, at each bout, the visual patterns were rotated by the amount specified by the bout bias multiplied by the closed-loop gain, on top of the exafferent stimulus rotation. Reafferent rotational velocity at each bout followed the shape of a decaying exponential with a time constant of 50 ms. In each experiment, fish received either only forward translational or rotational feedback, but never both.

### Behavioral Data Analysis

Behaviors of the fish were quantified in terms of the forward swim velocity or bout bias estimated online and downsampled to 20 Hz. Turning angular velocities were smoothed within 1 s rolling window when being plotted against time. These measurements were averaged across trials with the same stimulus conditions, averaged over time, correlated with or regressed by stimulus parameters, as detailed in the corresponding figure captions.

In the noise correlation experiments (Figure 4, Figure S4), forward or angular velocity of the stimulus *v*(*t*) was determined as *v*(*t*) = *v*_base_ × *s*(*t*) where *v*_base_ = 15 mm/s for the forward OMR experiment and *v*_base_ = 60 °/s in the rotational OMR experiment, and *s*(*t*) = −1 or +1. *s*(*t*) flipped its sign following a 0.5 Hz point Poisson process. The bout-triggered average (BTA) was calculated as

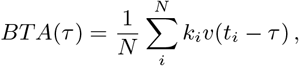

where *N* is the number of bouts, *k*_*i*_ is either peak swim speed (for forward OMR) or bias (for rotational OMR) of the i-th bout, and *t*_*i*_ is the onset timing of the i-th bout. For the forward OMR experiment, bouts with the bias higher than 1 rad were excluded from the analysis. For the both experiments, fish whose bout frequency was below 0.05 Hz was excluded, as the small number of bouts made it difficult to estimate the BTA. To quantify the timescales of the computed BTA stimuli, either a decaying exponential *ŷ* (*t*) = *y*_0_*e*^*t/τ*^ (for forward OMR) or a logistic function *ŷ* (*t*) = *y*_0_*/*(1 + *e*^(*t−t*0)*/τ*^) (for rotational OMR) were fit to the BTA curve from individual fish, as well as fish-averaged BTA curve. The logistic fit to the rotational BTA was chosen ad hoc as the sigmoid shape of the curve led to poor exponential fitting.

### Computational Modeling

The conditional leak model (Figure 5) was designed to reconcile the different timescales observed in the reverse correlation and priming-delay experiments (Figure 4). The model integrator unit activity *r*(*t*) followed the first-order dynamical equation

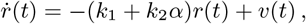

where *v*(*t*) is the velocity of the optic flow (i.e. model input), *k*_1_, *k*_2_ are constants, and *α* = 0 if |*v*(*t*) | *> v*_thresh_ otherwise *α* = 1. The time constant of the system becomes *τ*_1_ = 1*/k*_1_ if *α* = 0 (i.e. in the presence of optic flow) and *τ*_2_ = 1*/*(*k*_1_ + *k*_2_) if *α* = 1 (i.e. in the absence of optic flow). In Figure 5, we set *k*_1_, *k*_2_ such that *τ*_1_ =2.7 s and *τ*_2_ = 15 s, as well as *v*_thresh_ = 1.0 mm/s. The model unit activity *r*(*t*) was converted to the bout rate *λ*_bout_(*t*) = *λ*_base_ + (*λ*_max_ − *λ*_base_)*/*(1 + *e*^*−a*(*r*(*t*)*−b*)^), where *λ*_base_ = 0.05 Hz, *λ*_max_ = 2.0 Hz, *a* = 0.27, *b* = 15. For the reverse correlation simulation, bouts were stochastically simulated as a point Poisson process with a time varying rate *λ*_bout_(*t*). The simulation was performed at 10 Hz.

The saturation model (Figure S7A) and the normalization model (Figure 7E) were designed to explain the multiple timescales of the OMR hysteresis (Figure 4, Figure 5) based on the responses of IO neurons (Figure 7). Here, the stimulus velocity *v*(*t*) is first half-wave rectified:

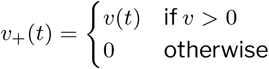

and

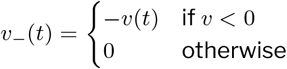

Next, the rectified velocities are separately integrated following the dynamical equation

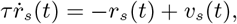

where *τ* = 15 s and *s ∈*{+,−}. In the saturation model, the OMR drive was computed as

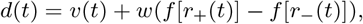

where *f* [*x*] = 1*/*(1 + *e*^*−βx*^), *w* = 5, *β* = 6. In the normalization model, the OMR drive was calculated as

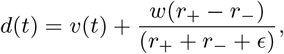

where *w* = 5 and *ϵ* = 0.01. The bout rate *λ*_bout_(*t*) was then calculated in both models as

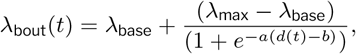

where *λ*_base_ = 0.05 Hz, *λ*_max_ = 1.0 Hz, *a* = 0.2, *b* = 15. The simulation was performed at 10 Hz.

### Light-sheet Microscopy

The custom-build light-sheet microscope used for the imaging experiments (Figure 6, Figure S6) was previously described in detail [12, 18]. The larvae were embedded in 2.0% agarose in a custom-built rectangular chamber, and submerged in the water from the fish facility. The chamber had glass-covered windows on the front and the side to let the excitation beams in. The agarose in front of the fish was removed such that it does not scatter the excitation beams. The agarose around the tail was also removed so that the fish can move its tail. The tail movements were monitored in the same way as in the behavioral experiments. The visual stimuli were also provided in the same way as in the behavioral experiments, except that the projector outputs were filtered with a long-pass filter (Kodak Wratten No.25) to allow imaging of green fluorescence. Each fish waited at least 1 hour after the embedding procedure before entering the experiments.

The excitation was provided by a 488 nm laser source (Cobolt), which was split between two orthogonal excitation arms. The two orthogonal beams allowed us to excite cells between the eyes and posterior to the eyes without exposing the eyes. Each arm consisted of a pair of galvanometric mirrors (Thorlabs) to scan the beam vertically and horizontally, a 2x beam expander, and a low numerical aperture air objective (Olympus) that focused the beams on the brain. The fast horizontal scanning of the laser created sheets of light. The emitted fluorescence was filtered with a 525:50 band pass filter and collected with a CMOS camera (Hamamatsu), through a 20x water immersion objective (Olympus). The collection objective was equipped with a piezo drive and moved in synchrony with the sawtooth-patterned vertical scanning of the light sheets, achieving the volumetric imaging. The resultant spatio-temporal resolution of the data was around 15 x 0.6 x 0.6 microns per voxel at 2 Hz. The control of the microscope was provided by a custom written software [42].

### Two-photon Microscopy

For the two-photon microscopy experiments (Figure 7), animals were embedded in a 35 mm petri dish in 2.0% agarose, in the same way as in the behavioral experiments. The visual stimuli were presented from below, and long-pass filtered in the same way as in the light-sheet imaging experiment. The excitation was provided by a femtosecond pulsed laser at the wavelength of 920 nm (Spark ALCOR 920-2) through a high NA, 20x water-immersion objective (Olympus). The average power at the sample was approximately 10 mW. The emitted fluorescence was filtered with a 750 nm short pass and a series of two band pass filters (535:40, 525:39). The scanning head consisted of a horizontally scanning resonant mirror (12 kHz) and a vertically scanning galvanometric mirror, controlled by a custom-written software through a National Instruments FPGA. The pixels were sampled at 20 MHz and averaged 8-fold, resulting in the 5 Hz frame rate with the image size of approximately 500 x 600 pixels and the resolution of about 0.2 micron per pixel.

### Stimulus Protocols for Imaging Experiments

At the beginning of each light-sheet imaging experiment (Figure 6), the pink noise pattern moving in the four directions (forward, backward, left, right) at 15 mm/s were presented in an open loop. Each movement lasted 5 seconds with 5 seconds of interleaves, and each direction was repeated 5 times. Next, the fish observed a continuous forward movement of the pink noise pattern at 5 mm/s for 30 minutes. During this period, swim bouts of the fish triggered visual feedback with a fixed velocity profile in a stochastic manner (superimposed on the constant flow). The feedback velocity was defined as

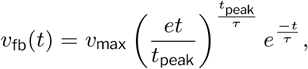

where *t* is the time since the bout onset. Here, *v*_fb_(*t*_peak_) = *v*_max_ and *τ* controls the fall-off. The parameters were set to mimic the vigor profile of a typical forward bout during the OMR (*v*_max_ = 50 mm/s, *t*_*peak*_ = 50 ms, *τ* = 100 ms). To dissociate the effects of the motor activity and visual feedback, for 50% of the bouts, visual feedback was not provided. In addition, pulses of backward flows with the same profile as described above were stochastically presented in the absence of bouts at the mean rate of 0.5 Hz, if fish made no bouts for more than 3 seconds.

The two-photon imaging experiment on the IO neurons (Figure 7) also started with the repeated presentation of the pink noise pattern moving in four directions as in the light-sheet imaging, but this portion of the data was not analyzed. The remaining of the experiment consisted of repetitions of 22 s long trials, which were all in an open-loop. At the beginning of each trial, a self-repeating pink noise pattern was reset to a random position. After an initial wait of 5 seconds, the pattern moved forward or backward for 2 seconds (“priming pulse”). In a subset of trials, immediately following the priming pulse, the pattern oscillated back and forth at 0.5 Hz with the peak speed of 5 mm/s. For the rest of the trial, the pattern remained stationary. Trials with and without oscillations were respectively repeated for 24 and 12 times for each priming pulse direction.

### Imaging Data Analysis

We used the suite2p package [43] to rigidly align the frames and to extract regions of interests (ROIs) from the raw calcium imaging movies, for both light-sheet and two-photon data. Briefly, suite2p iteratively aligns the movie to randomly picked reference frames using phase-correlation. To find neuronal somas, suite2p performs singular value decomposition on the movie and looks for peaks in the spatial singular vectors after smoothing. For the light-sheet imaging data, the aligned, time-averaged volumetric images of each brain were then registered to the MapZBrain atlas reference brain [44] using ANTsPy [45] in two steps: an affine transformation based on manually defined key-points, and a non-rigid, diffeomorphic registration. The coordinates of the ROIs in the MapZBrain space is calculated using this registration, and their affiliation to different brain regions were determined using anatomical masks provided on the MapZBrain. For the two-photon imaging data on the IO neurons, manual rectangular masks were drawn around the IO, according to which the ROI coordinates were normalized.

For the experiment in (Figure 6), we first identified direction selective visual neurons as follows: First, within the initial section during which translational motion was presented, the fluorescence time trace of each ROI *F* (*t*) was normalized into Δ*F/F* = (*F* (*t*) −*F*_0_)*/F*_0_, where the baseline *F*_0_ was defined to be the 1st percentile value of the raw fluorescence during this period. Next, the Pearson correlations of the normalized fluorescence between every pair among the five repetitions of the translational stimuli were calculated and averaged. Only ROIs with the averaged correlation above 0.4 are considered to be reliable visual neurons. Third, direction selectivity index (DSI) was calculated as 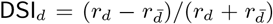 where *r*_*d*_ is the time- and repetition-averaged normalized fluorescence during the presentation of motion in direction *d* ∈ {left, right, forward, backward}, and 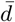 is the opposite direction of *d*. We considered an ROI to be selective for direction *d* only if DSI_*d*_ *>* 0.4 as well as *r*_*d*_ ≥*r*_*D*_ for all *D* left, right, forward, backward. Lastly, we calculated the triggered averages of the fluorescence for each of the three events (i.e., bouts without feedback, bouts with feedback, and backward flows without bout). Here, the raw fluorescence was converted to Δ*F/F*, using the average fluorescence during the 3 seconds preceding the onset of each event as *F*_0_. The triggered averages were then averaged across ROIs and fish (for visualizations of response dynamics), as well as across ROIs and over time within the 3 second window after the event onset within each fish (for statistical comparisons).

The two-photon imaging data on the IO (Figure 7) was first converted into Δ*F/F* for each ROI and each ROI, using the average fluorescence during the initial 5 second waiting period as *F*_0_. For each ROI, Δ*F/F* was averaged over time within a 2-s window during the priming phase (“early activity”), as well as within the last 9 seconds of the trial (“late activity”). Next, repetitions of the trials without the oscillatory stimuli were split into two halves (i.e., training and test sets). The training set priming response difference Δ*μ*_early_ = *μ*_fe_ −*μ*_be_, where *μ* indicates the mean forward early (fe) and backward late (be) activities, were calculated to examine the general topographic pattern of the direction selectivity in the IO (Figure 7c). To select ROIs that exhibited memory-like activity, for each ROI, Z-scored difference

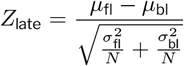

was calculated where *μ* and *σ*^2^ respectively indicate the mean and variance of the forward late (fl) and backward late (bl) activities, and *N* is the number of repetitions (i.e. 12). ROIs with |*Z*_late_| *>* 2 and congruently signed *Z*_late_ and Δ*μ*_early_ were selected as “memory cells”. The activities of memory cells were averaged across trials and ROIs within each fish for each condition (only using the test set for the conditions without oscillations), and then across fish, for the dynamics visualization (Figure 7E). The late activity was averaged within each fish to make comparisons across stimulus conditions (Figure 7F, G).

### Statistics

For comparisons of time-averaged behaviors or neural responses within each fish across conditions or against 0, we used Wilcoxon signed-rank tests. For comparisons of instantaneous swim or turn velocities within each fish across conditions, we used paired t-tests for the computational simplicity (Figure 1D, Figure 3B, Figure S2E).

**Table S1.** Stimulus protocols used for the experiments.

| Figure | Description |
| --- | --- |
| 1 | Each trial consisted of 5 seconds of "pre-wait", 3 seconds of "priming", 0 or 5 seconds of "wait", and 10 seconds of "probe" phases. At the beginning of each trial, a self-repeating pink noise pattern was reset to a random position. During the priming phase, the pattern moved at 5 mm/s in an open-loop, either forward or backward, with optional directional switching in the middle. During the probe, the pattern moved forward at 5 mm/s in a closed loop with the gain of 1. The pre-wait and wait phases were in an open-loop, and the pattern remained stationary. Each trial type was repeated 20 times. |
| 2 | The overall structure was the same as in Figure 1, but the priming and wait phases were respectively 5 s and 2 s long. During the priming and probe phases, the pattern always moved forward at 5 mm/s. The priming, but not probe, phase was in a closed-loop, with the gain of 0.5, 1.0, or 1.5. Each trial type was repeated 40 times. |
| S2A-C | The experiment consisted of 45 s long trials during which the pink noise pattern rotated at angular velocities of either 7.5, 15, 30, 60, or 120 °/s in either direction. The stimulation was in a closed-loop with a low gain of 0.3. Each trial type was repeated 8 times. |
| S2D, E | Each trial consisted of 10 s "pre-wait", 3 s "priming", 0 or 5 s "wait", and 20 s "probe" phases. Only the probe phase was in a closed loop, with the gain of 1. A pink noise pattern rotated at 30 °/s in either direction or stayed stationary in the priming phase, and rotated at 30 °/s in the probe phase in either direction. Each trial type was repeated 8 times. |
| 3 | The temporal structure of each trial was identical to that of Figure 2. During the priming phase, the pattern rotated at 30 °/s in either direction, with the closed-loop gain of 0.5 or 1.5. In the probe phase, the pattern rotated at 30 °/s in either direction. Each trial type was repeated 20 times. |
| 4A-D | A pink noise pattern moved forward or backward at 15 mm/s in an open-loop. The direction switched at random, following a Poisson point process with the rate of 0.5 Hz. The experiment lasted 1 hour. |
| S4A-D | A pink noise pattern rotated at 60 °/s in an open-loop. The direction switched at random, following a Poisson point process with the rate of 0.5 Hz. The experiment lasted 1 hour. |
| 4E-H | The overall structure was identical to Figure 1, but the duration of the wait phase was either 1, 2, 4, 8, or 16 seconds. The direction of flow during the priming phase was either forward or backward throughout, without switching. Each trial type was repeated 20 times. |
| 5F-I | The overall structure was identical to Figure 4E-H. In the half of the trials, the pattern sinusoidally oscillated back and forth with the frequency of 0.5 Hz and the peak amplitude of 5 mm/s during the open-loop wait phase, instead of remaining stationary. The duration of the wait phase was 2, 6, or 10 seconds. Each trial type was repeated 12 times. |
| S5 | The overall structure was identical to Figure 5F-I, but the wait phase duration was fixed at 6 seconds. A plaid pattern composed of two orthogonal (axial and lateral) sinusoids with the period of 10 mm was used as a visual pattern. During the wait phase, the pattern either stayed stationary or was modulated in following four different ways: (1) Standing wave: The axial component of the plaid oscillated as in a standing wave at 0.5 Hz. (2) Axial oscillation: The axial component of the plaid traveled back and forth sinusoidally, with a frequency of 0.5 Hz and a peak speed of 5 mm/s. (3) Lateral oscillation: The same as the axial oscillation, but the lateral component moved. (4) Luminance oscillation: Dynamic luminance offset was added to the whole pattern, which sinusoidally oscillated at 0.5 Hz, with the peaks of +/- 50% maximum luminance. Each trial type was repeated 15 times. |

## Supplementary figures

Seven supplementary supplementary figures follow.

**Figure S1:**
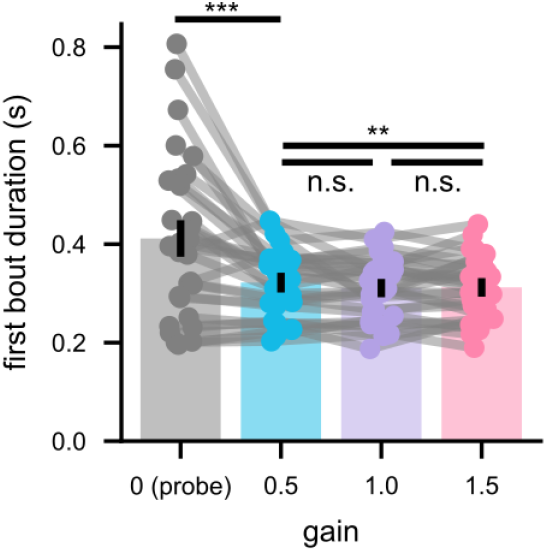
The fish are aware of gain manipulations. The average duration of the first bouts in the open-loop probe phase as well as the priming phase with different gains. As reported previously[12, 15], fish made significantly longer bouts in the lower gain conditions. The bars indicate the average across fish, and the dots individual fish. The data from identical fish are connected. N = 30 fish. P-values are from signed-rank tests. n.s.: p > 0.05; **: p < 0.01; ***: p < 0.0001.

**Figure S2:**
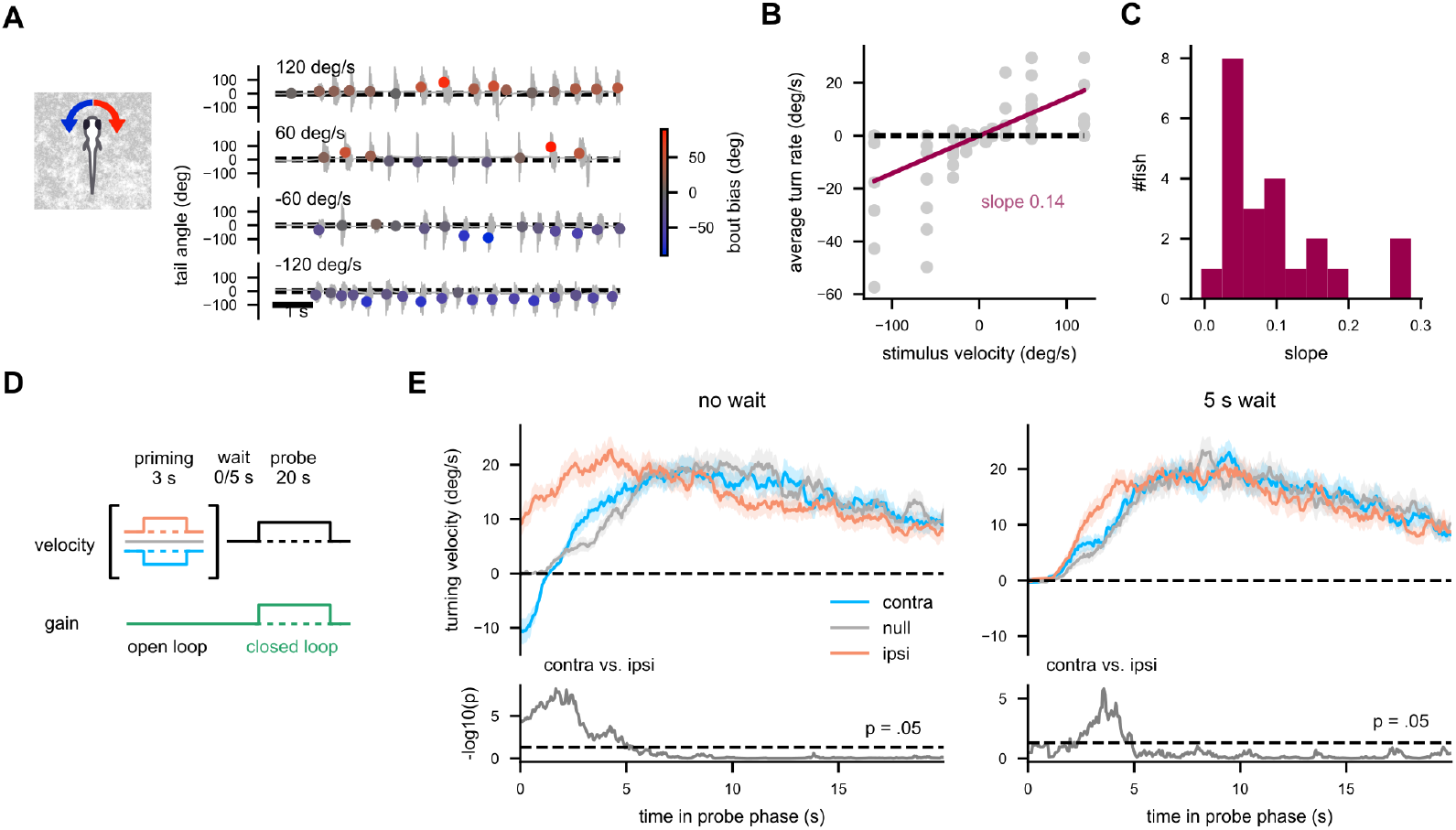
Rotational OMR in larval zebrafish and its history dependence. (A) Example tail traces of fish observing rotational stimuli with four different angular velocities. The dots indicate swim bouts, with y-position and color representing the bias of the bout. The fish tries to turn in the same direction as the stimulus, a classic phenomenology of the rotational OMR. (B) Turning velocity of the same single fish as in (A), averaged within each 45-s long trial (gray dots), with a linear fit by the stimulus velocity (red). (C) The histogram of the slope of the linear fit of fish’s average turn velocity by the stimulus angular velocity, as in (B). While most fish exhibited robust rotational OMR, the gain was generally low. (D) A schematic of the priming-probe assay for rotational OMR, similar to Figure 1B. (E) (top) The average angular velocity of the fish sorted by priming direction, and (bottom) p-values between ipsi- and contra-priming conditions (from paired t-tests). Conditions with and without a 5-second delay are plotted separately. Traces were folded and averaged such that the probe direction was positive. The angular velocity of the fish remained significantly different by priming directions until around 5 seconds into the probe phase. The solid lines and the shaded areas respectively indicate the mean across fish and its standard error. (B, C) N = 22 fish. (E) N = 29 fish.

**Figure S3:**
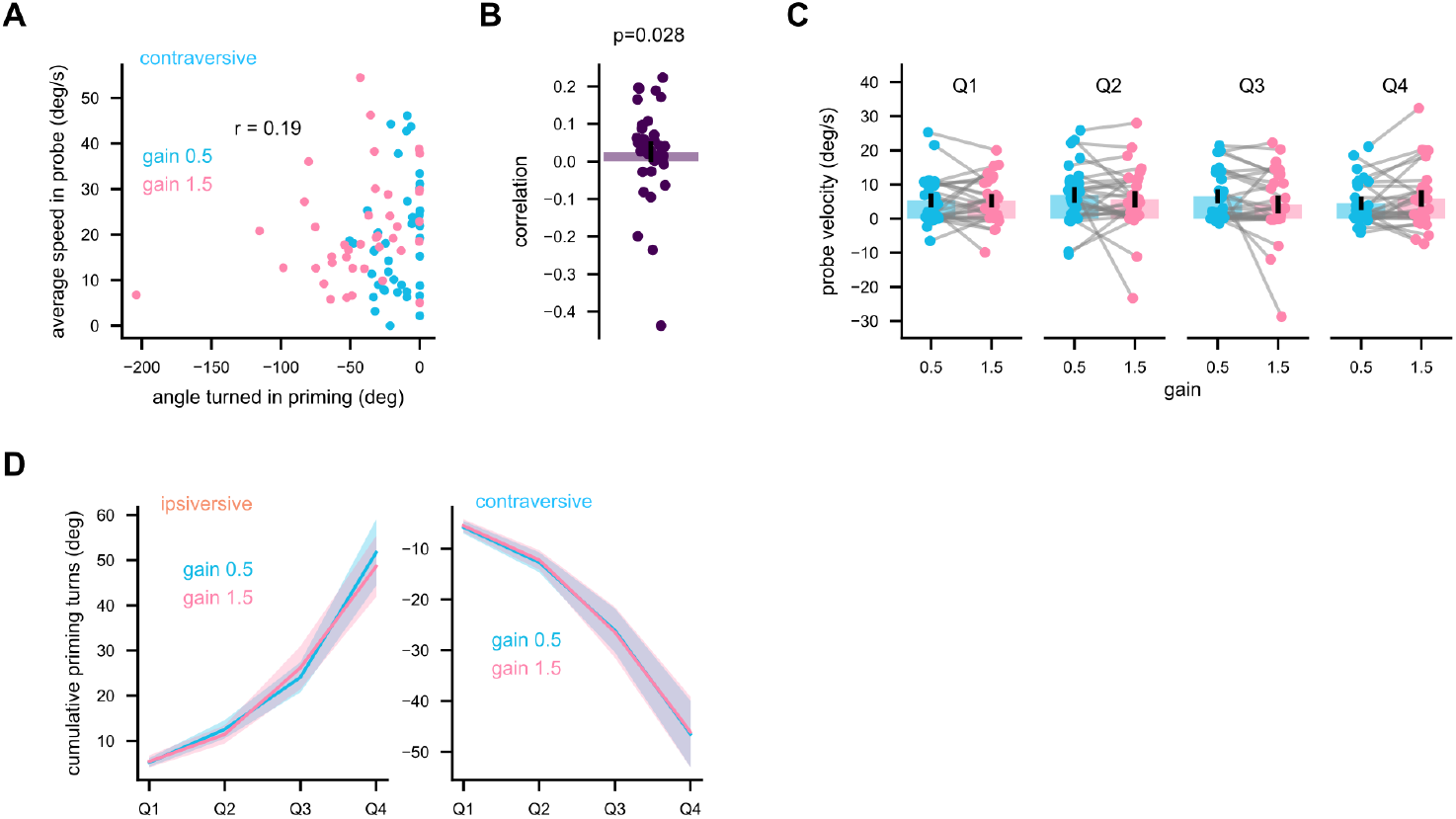
Rotational OMR with closed-loop, contra-directional priming. (A) The data from an example fish (same as in Figure 3C) in the contra-directional priming conditions. The turning velocities during the probe period (assuming 1.0 gain) are plotted against the angle turned in the priming period. Each dot corresponds to a trial. (B) The Pearson correlation between the probe speed and priming swim distance from all fish. The bar indicates the average across fish, and the dots individual fish. N = 36 fish. On average, less contra-directional swimming in the priming phase led to more OMR swimming in the probe phase, although the effect was weak and variable. This trend can be explained neither by the general alertness nor the homeostatic hypothesis. A likely source of this unexpected positive correlation is the tendency of fish to make consecutive turns in the same directions regardless of the stimuli[11]. (C) The trial-averaged probe turning velocities stratified into quartiles based on the cumulative tail angles during the priming phase for the contra-directional priming conditions. The data from identical fish are connected. No quartile showed significant effects of the gains (p > 0.05). N = 27, 29, 29, 28 for Q1, 2, 3, 4. (D) The cumulative tail angles during the priming phase in each quartile, used to stratify the probe speed in Figure 3E, Figure S3C. The amounts of turning effort fish made during the priming phase were matched well across the gain condition within each quartile. The solid line and the shaded areas indicate the mean across fish and its standard error. The p-values are from signed-rank tests.

**Figure S4:**
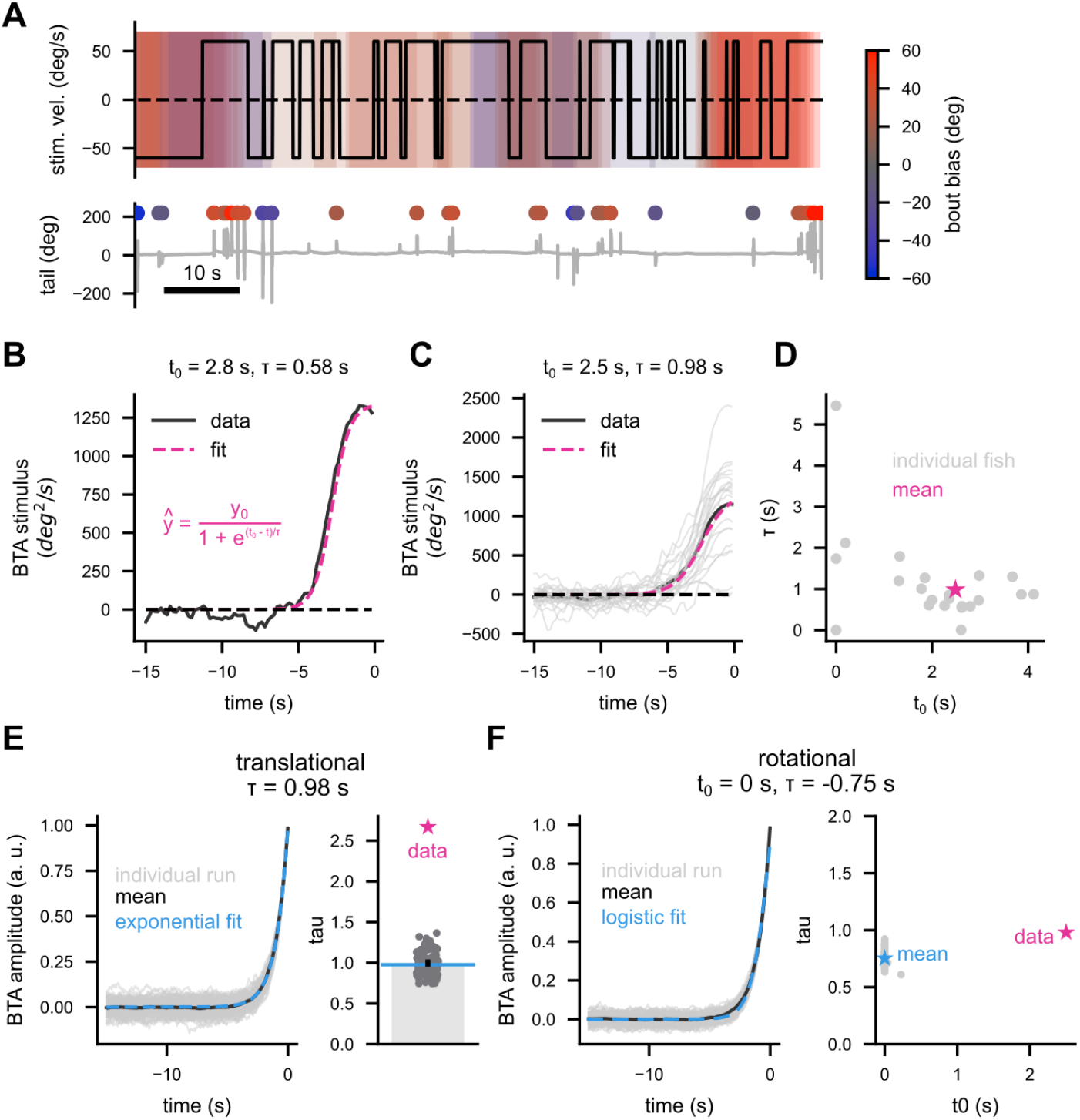
Additional characterizations of the history dependent OMR with the reverse correlation method. (A) Example traces of (top) stimulus velocity and (bottom) tail angles during the rotational reverse correlation experiment, similar to Figure 4A. (B) The bout-triggered average (BTA) stimulus angular velocity from a single fish (same as (A)) (black). A logistic function, as opposed to an exponential, was fit to the data (magenta), because the concavity around t = 0 led to poor exponential fits. The parameter t_0_ dictates when the bout triggered average reaches the half-maximum, and τ describes the steepness of the decay. (C) The population averaged BTA angular velocity (black), alongside the individual data (gray) and the logistic fit (magenta). (D) The two parameters of the logistic fit, plotted against each other for each fish and the population mean. The fits revealed that the rotational OMR has a time constant of about a couple seconds, similar to the forward translational OMR. N = 22 fish. (E, F) Simulated BTA for memory-less (E) translational and (F) rotational OMR. For both cases, we simulated an hour-long trace of velocities switching directions at 0.5 Hz. For the translational case, simulated forward bouts were generated following a memory-less Poisson point process with the rate of 1.0 Hz as long as the stimulus direction was positive. For the rotational case, simulated turning bouts were generated stochastically throughout, whose sign was identical to the direction of the stimulus at the moment. The BTAs (plotted over time in the left panels) were calculated in the same way as in the experiments, and exponential or logistic functions (cyan) were fit to the BTA of the individual simulation run (gray) or the mean BTA across 100 repetitions (black). The right panels show the time constant parameters for individual simulation runs (gray), fit parameters on the mean (cyan), as well as the parameters from the experimental data (magenta). In both cases, the time constants of the simulated BTA were around 1.0 s, well below those observed in the experiments.

**Figure S5:**
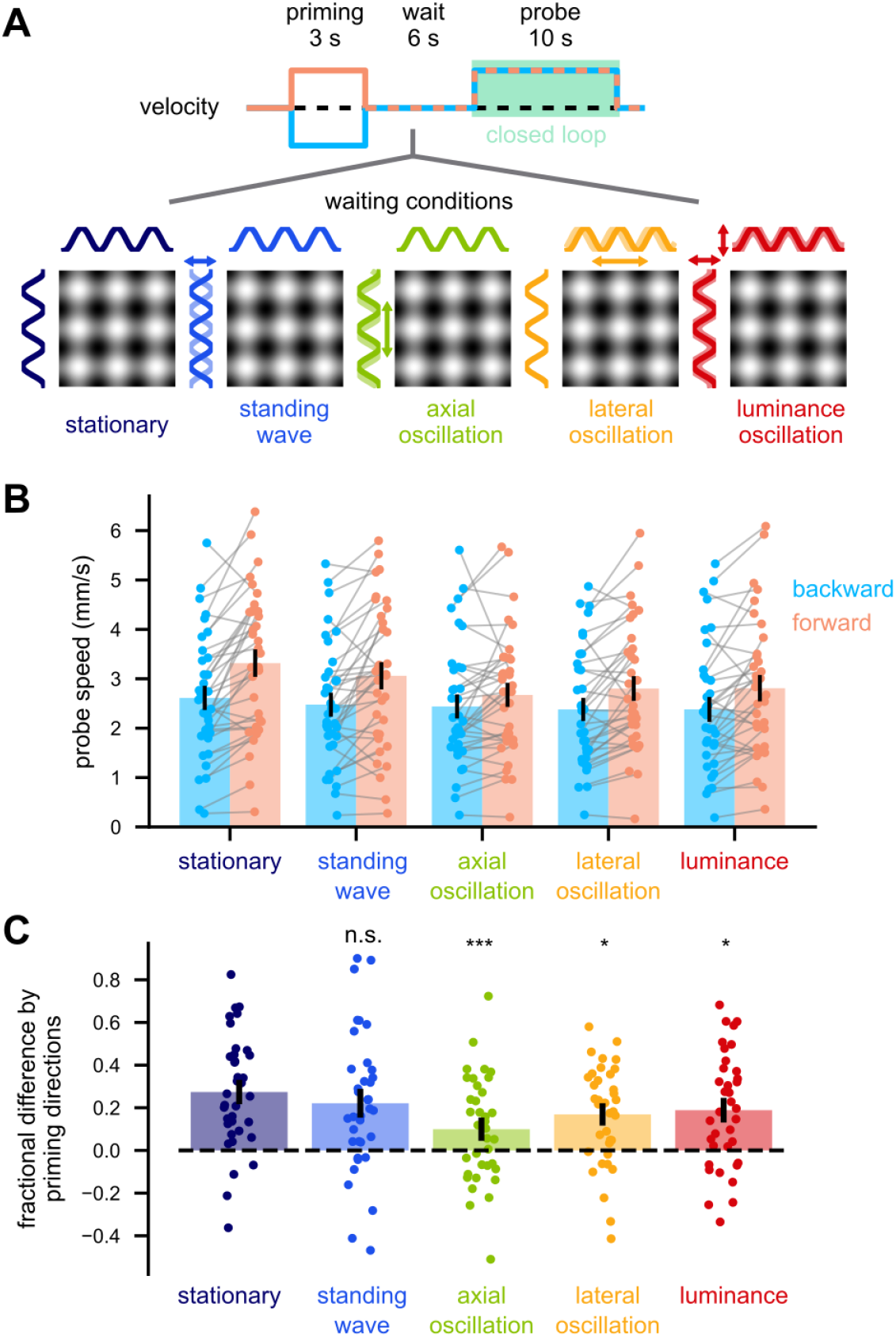
Forward and backward translation most strongly affects history dependent forward OMR. (A) A schematic of the protocol. The protocol structure with open-loop priming and closed-loop probe phases was similar to those in Figure 1, Figure 4, Figure 5. A plaid pattern with orthogonal sinusoids was presented instead of the pink noise pattern used in the other experiments. During the 6 s waiting period, the pattern remained stationary, or was modulated in four different ways as depicted in the panel. (B) The mean probe swimming speed by the priming directions and the waiting conditions. (C) Fractional difference in probe swimming speed by priming conditions, for each waiting condition. The “axial oscillation” condition resulted in the most significant decrease in optic flow memory compared to the stationary condition. Interestingly, the “standing wave” condition, which is a superimposition between forward and backward traveling waves, did not result in the reduction of optic flow memory. n. s.: p > .05; *: p < .05; ****: p < .0001 from signed-rank tests against the stationary condition.

**Figure S6:**
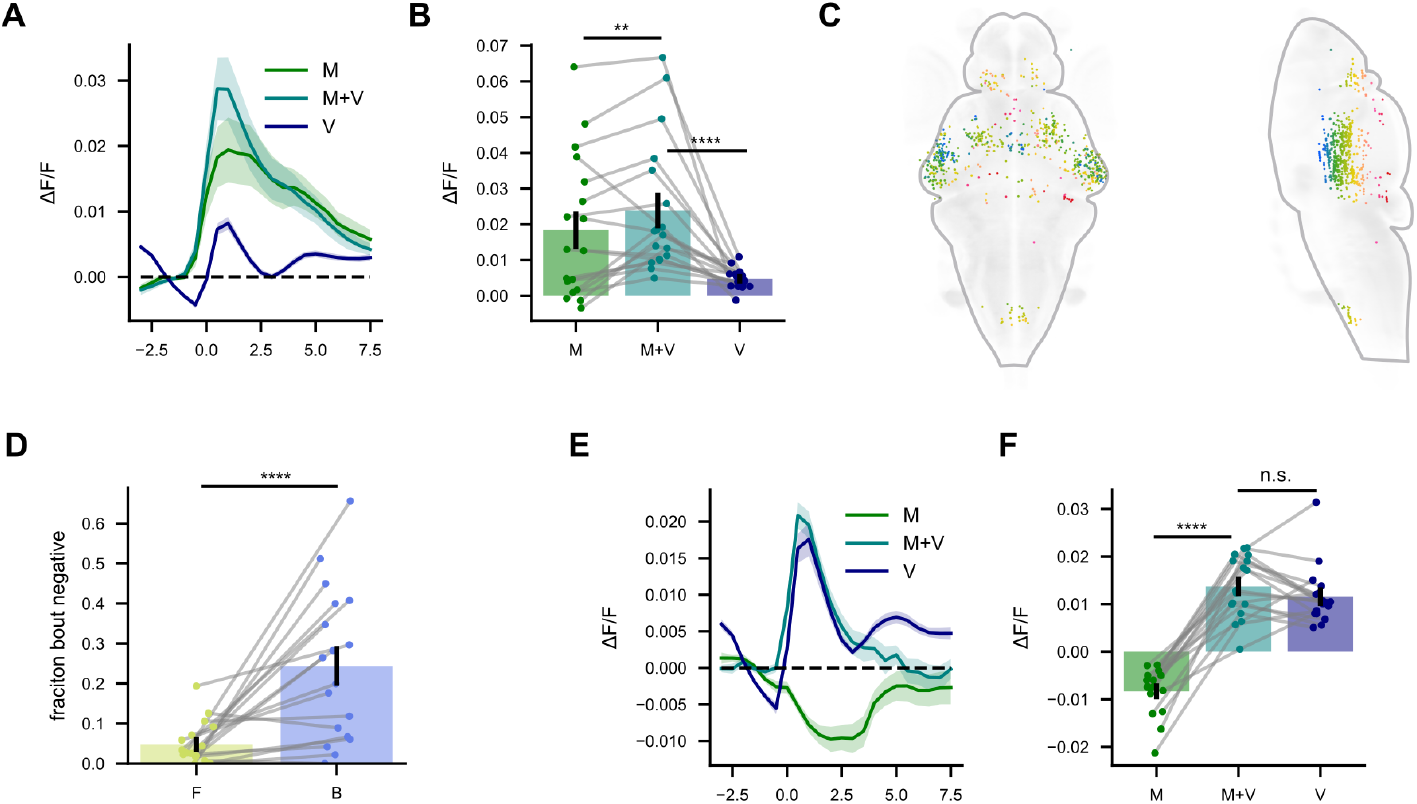
Motor-triggered activity in backward-tuned neurons. (A, B) Activity of backward-tuned neurons triggered by the three types of events (A) over time or (B) averaged across time, similar to Figure 6D, E. As expected from their direction selectivity, visual-only (V) events trigger positive activities. However, on average, these cells exhibited a positive motor triggered activity, which does not cancel the visually evoked activities. (C) The distribution of backward-tuned cells that exhibited negative (i.e., opposite to visual) response after a swim bout without visual feedback (i.e., M event). A majority of these backward-tuned, bout-negative cells were found in ventral optic tectum. (D) The fraction of bout-negative cells among the backward-tuned cells were significantly higher than that among the forward-tuned cells, suggesting some degree of directional specificity in efference copy signals delivered to optic flow tuned neurons. (E, F) Same as (A, B), but restricted to the bout-negative cells among backward tuned cells. Even among these cells, we did not observe significant cancellation of the visually triggered positive activity by the motor-triggered negative activity. This might conceivably because the fixed backward visual feedback fish received consistently exceeded the expected amount of feedback. Throughout, the solid lines and shaded regions represent the means across fish and their standard errors. Dots in the bar plots indicate individual fish, and the data from the same fish are connected. N = 18 fish. n.s.: not significant; *: p < .05, **: p < .01; ***: p < .001; ****: p < .0001 from signed-rank tests.

**Figure S7:**
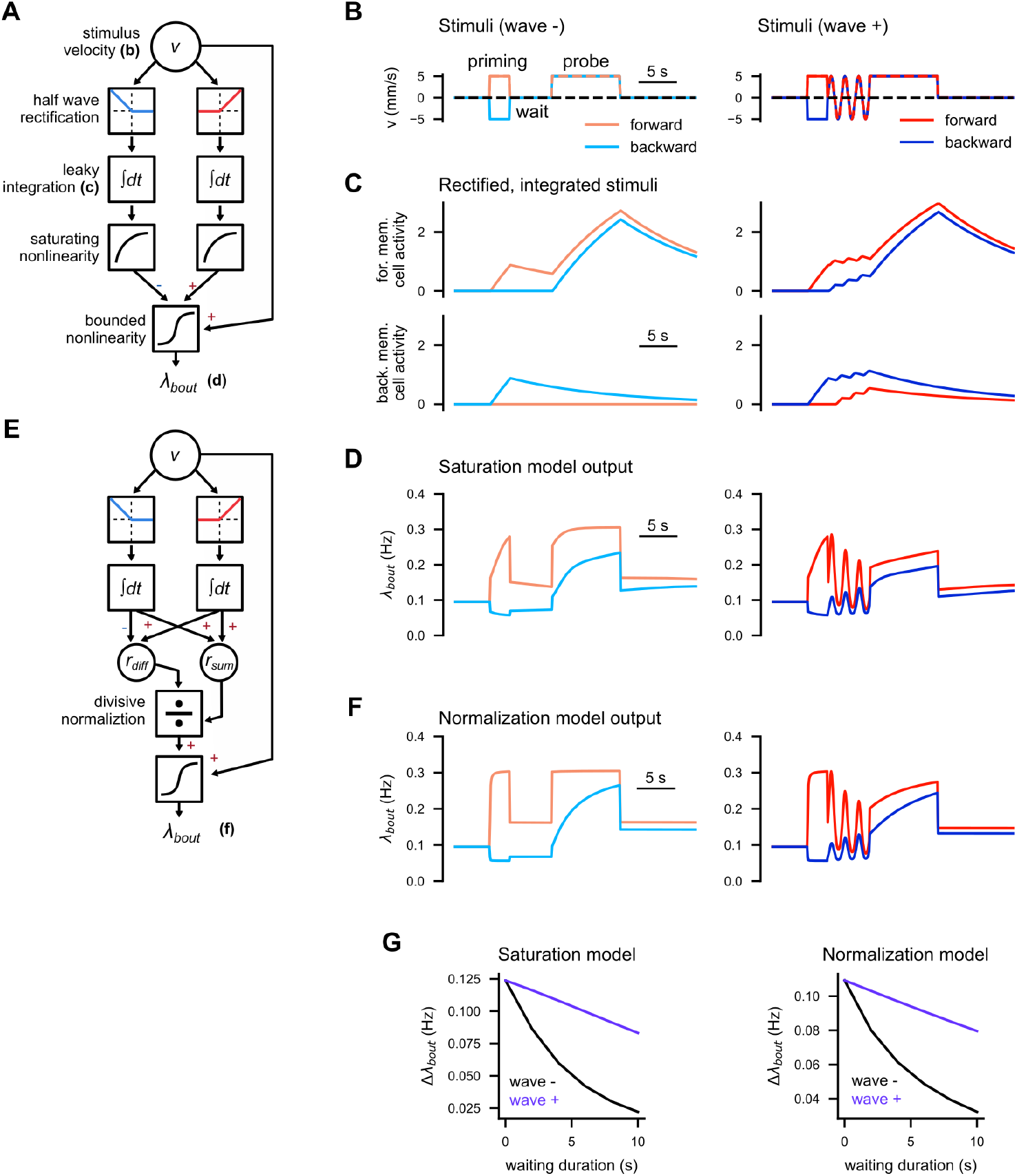
Simple models to connect the IO physiology to the behavior. Contrary to the naive expectation from the conditional leak model in Figure 5, the activity of IO memory cells was increased by oscillating stimuli regardless of their directional tuning (Figure 7E, F). To explain the faster forgetting caused by the oscillating stimuli (Figure 5H, I), one can simply assume a sublinearity between the memory cell activity and the bout frequency. (A) A schematic of a model featuring saturating nonlinearity (see Methods for the details). The stimulus velocity inputs are first half-wave rectified, and the forward and backward velocities are integrated separately in a leaky fashion (τ = 15 s). The integrated velocities each undergo a saturating nonlinearity, and then they are summed together with the stimulus velocity (with appropriate sign inversions). The combined signal then undergoes a bounded nonlinearity to generate the bout frequency λ_bout_. (B) The simulated stimulus time traces mimicking the priming-delay assay (Figure 5F), with or without the oscillation. (C) The rectified, integrated (top) forward and (bottom) backward velocities. See how they mimic IO responses in Figure 7E. (D) The time traces of λ_bout_ generated by the saturation model in (A). (E) A schematic of a model featuring divisive normalization. In this model, a ratio of the difference (r_diff_) and the sum (r_sum_) between the rectified, integrated forward and backward velocity is calculated, and is combined with the stimulus velocity to generate λ_bout_. (F) The λ_bout_ outputs of the normalization model in (E). (G) The difference in average λ_bout_ during the probe period between forward and backward priming conditions, as a function of the waiting duration, from (left) saturation and (right) normalization models. The both models replicate the shortening of optic flow memory by oscillating stimuli shown in Figure 5I.

## Notes

### Competing Interest Statement

The authors have declared no competing interest.

